# A new stegosaur (Dinosauria: Ornithischia) from the Upper Jurassic Qigu Formation of Xinjiang, China and a revision on Chinese stegosaurs phylogeny

**DOI:** 10.1101/2024.09.29.615678

**Authors:** Li Ning, Chen Guozhong, Octávio Mateus, Jiang Tao, Xie Yan, Li Daqing, You Hailu, Peng Guangzhao

## Abstract

Stegosaurs are a small but iconic clade of ornithischian dinosaurs. They and their sister taxa, the ankylosaurs, formed the clade Eurypoda which means ‘broad-footed’. Here, we describe a stegosaur from the Upper Jurassic Qigu Formation of Xinjiang, China, based on an associated partial skeleton that includes axial, pectoral girdle, pelvic girdle, limb and armor elements. It can be diagnosed as a new taxon, *Angustungui*, based on numerous autapomorphies. Some morphologies of *Angustungui* are more similar to the taxa from Europe, Africa and North America than to those from Asia. Our phylogenetic analysis recovers it as the sister taxon of *Loricatosaurus*. More importantly, the narrow and claw-shaped ungual of *Angustungui* proves that Eurypoda, at least stegosaur, has claw-shaped unguals. Besides, we revised the character scores for Chinese stegosaurs based on observations of the specimens.

## INTRODUCTION

Stegosaurs are known for having two parasagittal rows of hypertrophied dermal armour plates and/or spines extending from the neck to the tail (Galton and Upchurch 2004). They have been discovered on every continent except Antarctica and Australia. The earliest stegosaurs lived in the Middle Jurassic, such as *Huayangosaurus*, *Bashanosaurus*, *Baiyinosaurus*, *Loricatosaurus*, *Adratiklit*, *Thyreosaurus* and *Isaberrysaura* (Dai et al. 2022; Galton 1985; Li et al. 2024a; Maidment et al. 2020; Salgado et al. 2017; Sereno and Dong 1992; Zafaty et al. 2024). By the Late Jurassic, they achieved a global distribution, such as *Stegosaurus*, *Hesperosaurus* and *Alcovasaurus* from North America (Carpenter et al. 2001; Galton and Carpenter 2016; Maidment et al. 2015), *Dacentrurus* and *Miragaia* from Europe (Galton 1991; Mateus et al. 2009; Sánchez-Fenollosa et al. 2024), *Chungkingosaurus*, *Tuojiangosaurus*, *Gigantspinosaurus* and *Jiangjunosaurus* from Asia (Hao et al. 2018; Jia et al. 2007; Maidment and Wei 2006) and *Kentrosaurus* from Africa (Hennig 1915). However, their numbers declined during the Early Cretaceous, represented by *Paranthodon*, *Wuerhosaurus*, *Mongolostegus* and *Yanbeilong* (Dong 1990, 1993; Galton and Coombs 1981; Jia et al. 2024; Tumanova and Alifanov 2018).

Dinosaur fossils are abundant in the Xinjiang Uygur Autonomous Region of China. Since the first dinosaur from Xinjiang was discovered in 1930 (Young 1937), more than 20 dinosaur taxa have been discovered and named. Among these dinosaurs, stegosaurs include *Wuerhosaurus homheni* and *Jiangjunosaurus* (Dong 1990; Jia et al. 2007). Shanshan of the Turpan Basin is a famous location of fossil dinosaurs in Xinjiang, China. Two mamenchisaurid sauropods, *Hudiesaurus sinojapanorum* (Dong 1997) and *Xinjiangtitan shanshanensis* (Wu et al. 2013), and an allosaurid theropod, *Shanshanosaurus huoyanshanensis* (Dong 1977b), have been described. In 2016, a skeleton of stegosaur was discovered by Dr Li Daqing and his team from the Late Jurassic Qigu Formation at Qiketai, Shanshan County during field exploration. It is the first discovery of stegosaurian remains from the Turpan Basin of Xinjiang and is described in this paper. In addition, we revised the character scores for Chinese stegosaurs based on observations of the specimens.

### Geological setting

The material of the new taxon was found in Qiketai town, Shanshan County, Turpan City, Xinjiang Uygur Autonomous Region, China (Fig. 1A, B). The fossil site is located in the central part of the Turpan Basin, where the Jurassic strata are well exposed. The Qigu Formation is in conformable contact with the underlying Middle Jurassic and the overlying Cretaceous (Fig. 1D). This formation consists of fluvial-lacustrine deposits, with lithologies dominated by purple-red and grey-green siltstones, mudstones, sandstones, and sandy conglomerates. The Qigu Formation is divided into three members: the lower member consists of interbedded purplish-red and grey-green siltstone and mudstone; the middle member comprises interbedded purplish-red siltstone, mudstone and grey-green mudstone, siltstone; the upper member has a basal layer of brownish-red sandy conglomerate, followed by interbedded brownish-red and grey-green siltstone and mudstone (Zhang 2019). Stegosaur fossils were collected from the grey-green siltstone of the middle member of the Qigu Formation (Fig. 1C). The zircon U-Pb age of the lower part of the Qigu Formation is 151 Ma (Fang et al. 2015), which places the fossil beds in the Kimmeridgian-Tithonian age.

**Figure 1.**
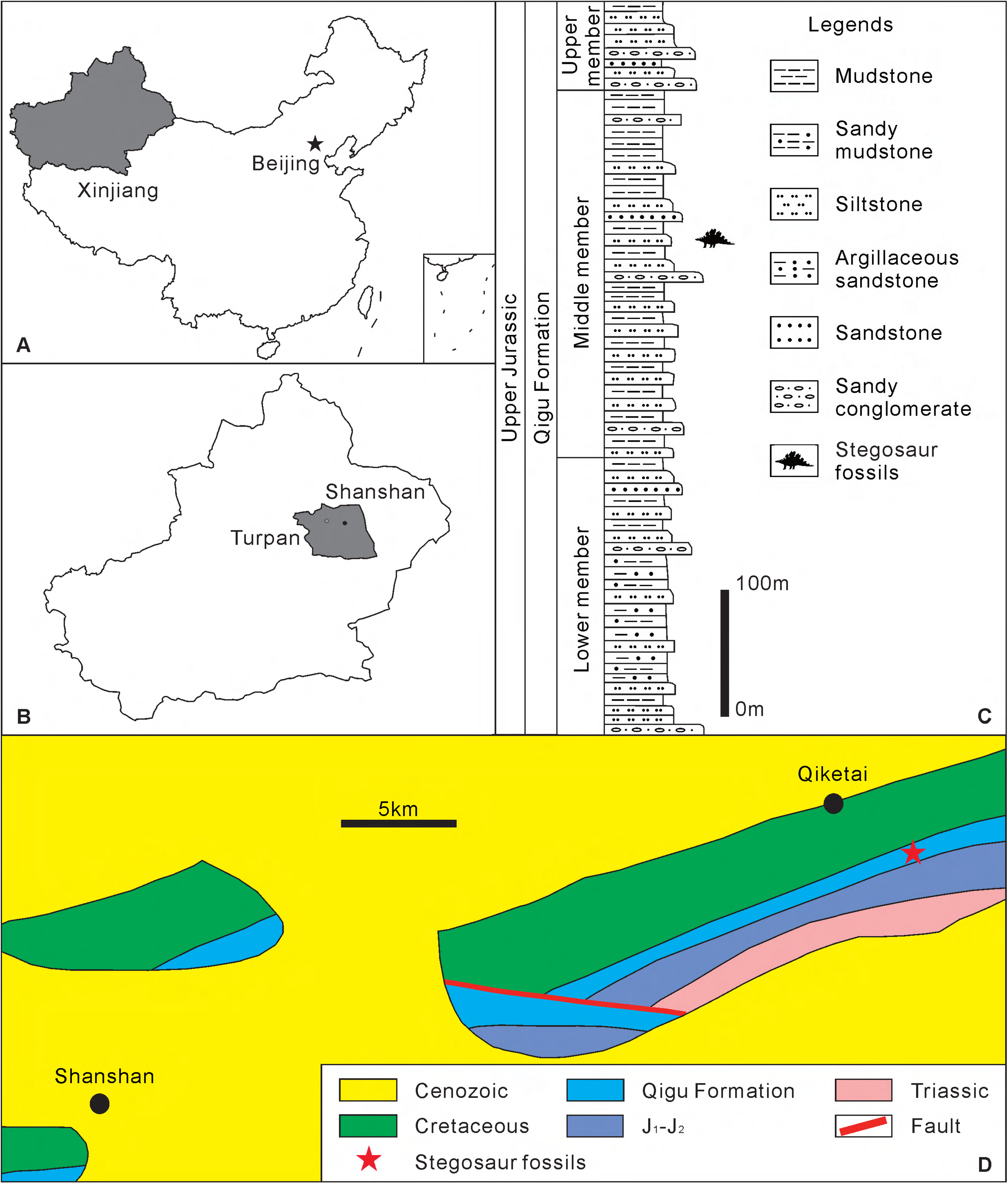
A, the map of China showing Xinjiang Uygur Autonomous Region. B, the map of Xinjiang Uygur Autonomous Region shows the Turpan City and Shanshan County. C, stratigraphic section of the Qigu Formation in the stegosaur locality, modified from Zhang Zhang 2019. D, Geological map of Qiketai town and fossil locality.

### Institutional abbreviations

BMNHR, The Natural History Museum, London, UK.

CQMNH, Chongqing Museum of Natural History, Chongqing Municipality, China. IVPP, Institute of Vertebrate Paleontology and Paleoanthropology, Chinese Academy of Sciences, Beijing, China.

MB, Museum für Naturkunde, Berlin, Germany.

SS, Shanshan Jurassic Museum, Shanshan, Turpan, Xinjiang Uygur Autonomous Region, China.

SXMG, Shanxi Museum of Geology, Taiyuan, Shanxi Province, China. ZDM, Zigong Dinosaur Museum, Zigong, Sichuan Province, China.

## MATERIALS AND METHODS

The specimen SS V16001 consists of a partial stegosaurian skeleton, including one cervical vertebra, ten dorsal vertebrae, several dorsal ribs, a synsacrum with one dorsosacral and four sacral vertebrae, seven caudal vertebrae, a right scapula, both ilia, both ischia, both pubes, a left femur, a phalanx, an ungual, a left parascapular spine and a dorsal plate. All the elements were associated or semi-articulated and belonged to one individual. During the field excavation, one free cervical vertebra and five free caudal vertebrae were collected alone and other bones were collected with a plaster wrap (Fig. 2). In addition, SS V16002 consists of a right scapula and a right coracoid, collected several meters away from SS V16001.

**Figure 2.**
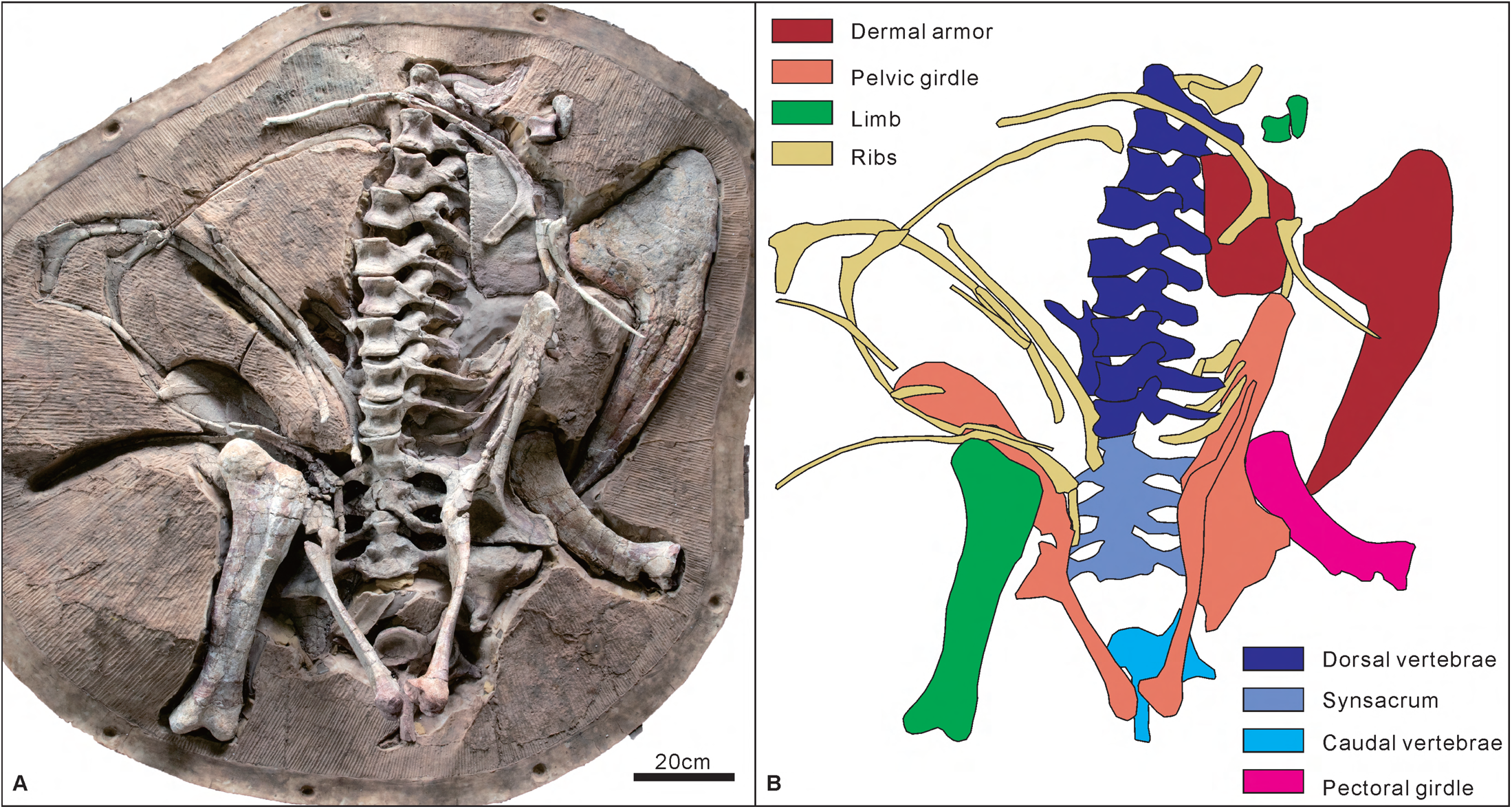
*Angustungui qiketaiensis* skeleton of the holotype (SS V16001): A, photograph as originally preserved. B, schematic drawing. Scale = 20 cm.

Raven and Maidment 2017 (2017) developed a stegosaur data matrix of 23 taxa scored for 115 characters, which was expanded upon in subsequent iterations (Dai et al. 2022; Li et al. 2024b; Maidment et al. 2020), with the version published by Li et al. (2024b) consisting of 27 taxa scored for 117 characters. We add the recently described early diverging thyreophoran *Yuxisaurus kopchicki* (Yao et al. 2022), stegosaur *Yanbeilong ultimus* (Jia et al. 2024) from China and *Thyreosaurus atlasicus* (Zafaty et al. 2024) from Morocco into the matrix. We also revised the mistakes in the character list and the character scores of the stegosaurs from China. The revised character scores are all from personal observations based on fossil specimens or previously published literature. To assess the phylogenetic relationships of *Angustungui*, we scored it for the modified version of the character-taxon matrix that consists of 31 taxa scored for 117 characters. The matrix was analysed in TNT v1.5 (Goloboff et al. 2008). *Pisanosaurus* was set as the outgroup. All continuous characters (1-25) and characters 108 and 109 were ordered. A New Technology search was performed using sectorial, ratchet, drift and tree fusing options and 10 random addition sequences. A second round of TBR branch-swapping was then performed on the trees held in Random Access Memory (RAM) to more fully explore treespace for additional most parsimonious trees (MPTs). Support for the relationships obtained was evaluated using Bremer support and bootstrap analysis (1000 replicates, traditional search).

## RESULTS

### Systematic palaeontology

> **Dinosauria** Owen 1842, 1842

> **Ornithischia** Seeley 1888, 1888

> **Thyreophora** Nopcsa 1915, 1915

> **Stegosauria** Marsh 1877, 1877

> **Stegosauridae** Marsh 1880, 1880

> *Angustungui qiketaiensis* gen. et sp. nov.

*Holotype*: SS V16001, a partial skeleton including one cervical vertebra, ten dorsal vertebrae, several dorsal ribs, a synsacrum with one dorsosacral and four sacral vertebrae, seven caudal vertebrae, a right scapula, both ilia, both ischia, both pubes, a left femur, a phalanx, an ungual, a left parascapular spine and a dorsal plate. Measurements of all elements of the holotype can be found in Supplementary 1.

*Paratype*: SS V16002 including a right scapula and a right coracoid. Measurements of all elements of the paratype can be found in Supplementary 1.

*Etymology*: *Angustungui*, after the Latin *angusti* (narrow) and *ungui* (claw), in reference to its narrow claw; *qiketaiensis* is derived from Qiketai (the town name of the type locality).

*Locality and horizon*: Qiketai town, Shanshan County, Turpan City, Xinjiang Uygur Autonomous Region, China. The materials are from the middle member of the Qigu Formation, Late Jurassic, assigned to the Kimmeridgian-Tithonian in age.

*Diagnosis*: *Angustungui* differs from all other stegosaurs by the presence of the following autapomorphies: (1) the anterior centroparapophyseal lamina (ACPL) of the dorsal vertebrae drawn into the anterolateral margin of the centrum; (2) in the ilium, the preacetabular process projects approximately parallel to the parasagittal plane in dorsal or ventral view and lies approximately horizontal in lateral view, and a ventromedial flange backs the acetabulum; (3) the ungual phalanges are claw-shaped; (4) the basal plate of the parascapular spine is subtriangular with a flat and blunt posterior margin in dorsal and ventral views, and the spine is strong such that the base of the spine length is more than one-third of the lateral margin of the basal plate length. Phylogenetic analysis recovered three apomorphies for *Angustungui*: ch. 9, 68 and 104.

### Description and comparisons

#### Axial skeleton

*Cervical vertebra*: One free posterior cervical vertebra is preserved, but the prezygapophyses, postzygapophyses, diapophyses and neural spine are missing (Fig. 3A-F). The centrum is longer anteroposteriorly than broad transversely, differing from *Dacentrurus* in that the centrum is wider than long (Galton 1985). The anterior and posterior articular surfaces are funnel-like concave. The posterior articular surface is more concave than the anterior articular surface, with a notochord depression in the centre (Fig. 3B), the same as *Jiangjunosaurus* (IVPP V14724). The ventral surface of the centrum is nearly flat (Fig. 3F), unlike *Stegosaurus* with a strong ventral keel (Maidment et al. 2015). The ventral keel of the centrum is also present in the posterior cervical vertebrae of *Jiangjunosaurus* (IVPP V14724). In lateral view, the ventral margin of the centrum is nearly straight. The lateral surfaces of the centrum are deeply concave (Fig. 3C, D), similar to *Dacentrurus* and *Loricatosaurus* (Galton 1985, 1991), but different from other stegosaurs such as *Huayangosaurus*, *Miragaia*, *Gigantspinosaurus*, *Stegosaurus* and *Jiangjunosaurus* (Hao et al. 2018; Jia et al. 2007; Maidment et al. 2006, 2015; Mateus et al. 2009; Zhou 1984). The parapophyses are located on the anterodorsal part of the centrum in lateral view and the dorsolateral margin in anterior or posterior view. The position of the parapophyses is more dorsal than in the seventh cervical vertebra of *Dacentrurus* and more similar to in the tenth cervical vertebra of *Dacentrurus* (Galton 1991), which indicates that it is a posterior cervical vertebra. Laterally, the neural arch is smooth and the neurocentral suture is visible, which indicates that this individual is a subadult. The base of the diapophysis is located at the base of the neural arch in the mid-length of the centrum (Fig. 3C). The neural canal has a subcircular outline in anterior and posterior views and is larger in the posterior view.

**Figure 3.**
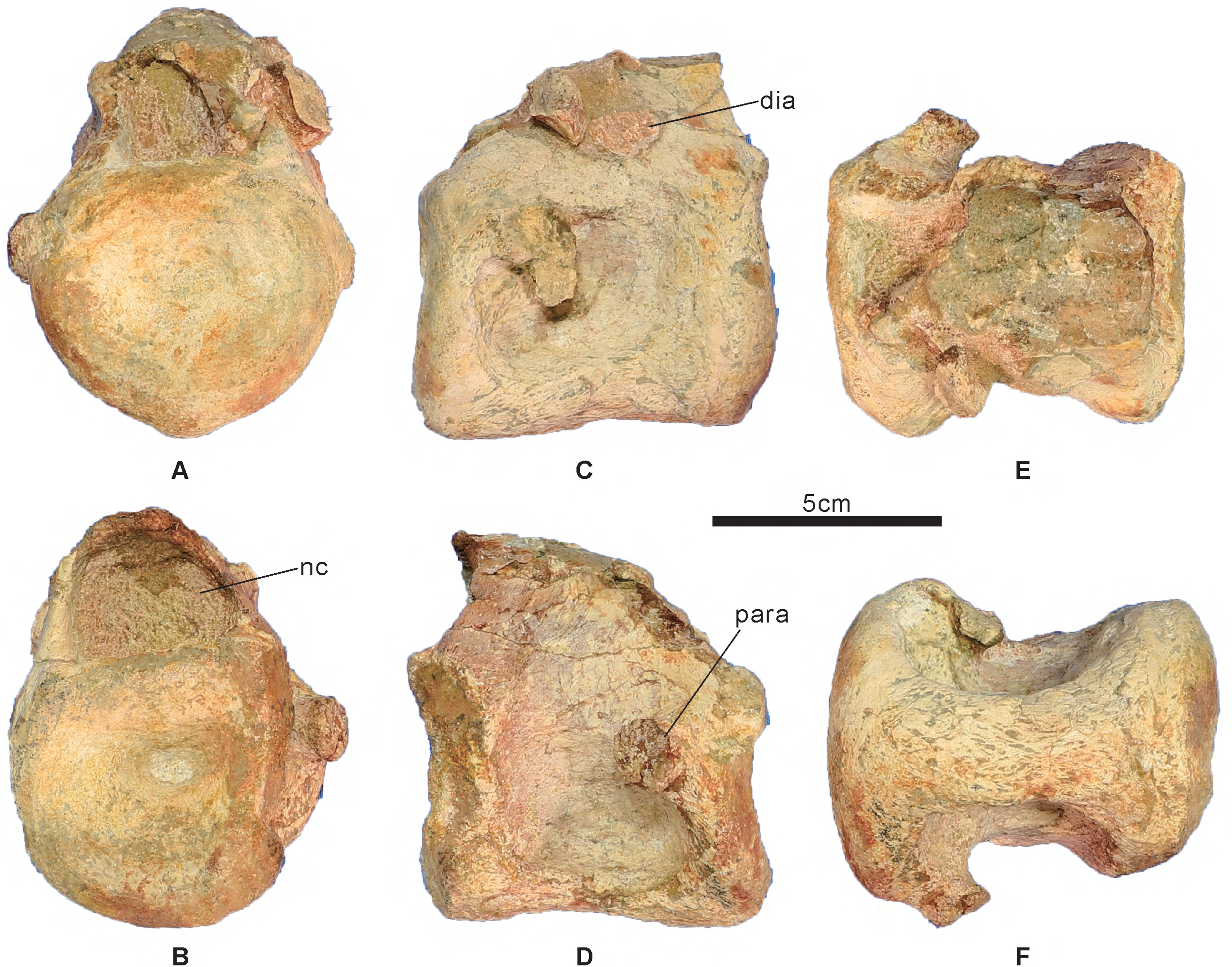
Cervical vertebra of *Angustungui qiketaiensis* (SS V 16001, holotype). A, anterior view. B, posterior view. C, left lateral view. D, right lateral view. E, dorsal view. F, ventral view. Abbreviations: dia, diapophysis; nc, neural canal; para, parapophysis. Scale = 5 cm.

*Dorsal vertebrae*: Ten nearly complete dorsal vertebrae are preserved, including one free anterior dorsal vertebra and nine articulated posteriormost dorsal vertebrae (Fig. 2, 4B-D). The last dorsal vertebra is articulated with the sacrum (Fig. 2). The centrum of the free dorsal vertebra and the first to seventh of the nine articulated dorsal vertebrae is longer anteroposteriorly than broad transversely and the centrum of the last two dorsal vertebrae is wider than long, which differs from *Adratiklit* (Maidment et al. 2020) and *Dacentrurus* (Galton 1985; Sánchez-Fenollosa et al. 2024) that all the centrum is wider than long. All the dorsal centra are amphicoelous, with a circular outline, similar to other stegosaurs such as *Stegosaurus*, *Hesperosaurus*, *Bashanosaurus*, *Yanbeilong* and *Baiyinosaurus* (Carpenter et al. 2001; Dai et al. 2022; Jia et al. 2024; Li et al. 2024a; Maidment et al. 2015). The preserved dorsal vertebrae 7 and 9 exhibit distinct ventral keel (Fig. 4D). The lateral surfaces are deeply concave anteroposteriorly, similar to the cervical vertebrae (Fig. 2A, 4A-B), but different from other stegosaurs except *Dacentrurus*, *Adratiklit* and *Thyreosaurus* (Galton 1985; Maidment et al. 2020; Zafaty et al. 2024). The top of the centrum merges with the base of the pedicels, and the neurocentral suture is also visible (Fig. 4B). The base of the pedicels is convex laterally relative to the deeply concave of the lateral surfaces of the centra (Fig. 4A). The pedicels are short and extend anterodorsally. In the dorsal vertebral series, the pedicels gradually increase in height. A ridge on the pedicel, the anterior centroparapophyseal lamina (ACPL), extends from the anteroventral corner of the parapophysis anteroventrally towards the anterolateral margin of the centrum (Fig. 4A). The ACPL is more developed in the posterior dorsal vertebrae. However, in *Stegosaurus* and *Bashanosaurus*, the ACPL of the dorsal vertebrae merges ventrally with the neural arch before reaching the neurocentral suture (Dai et al. 2022; Maidment et al. 2015) and it is drawn into anteriorly-projecting rugosities either side of the neural canal in *Adratiklit* (Maidment et al. 2020). In posterior view, there is a dorsoventrally medial keel on the pedicel (Fig. 4D). In the posterior view, the neural canal has a teardrop-shaped outline (Fig. 4D), different from the *Kentrosaurus* that has an extremely enlarged neural canal (Galton 1982). The parapophysis is situated anteroventral to the base of the diapophysis and poorly developed, different from *Bashanosaurus* where the elevation of the parapophysis is greater (Dai et al. 2022). The parapophysis has a sub-circular outline and is gently concave. The prezygapophyses extend anterodorsally. The articular facets of the prezygapophyses are flat and face dorsomedially, similar to most stegosaurs such as *Kentrosaurus*, *Hesperosaurus* and *Baiyinosaurus* (Carpenter et al. 2001; Galton 1982; Li et al. 2024a). There is a distinct dorsal process at the dorsolateral margin of the prezygapophysis (Fig. 4A), which is also present in *Dacentrurus* and *Adratiklit* (Maidment et al. 2020; Sánchez-Fenollosa et al. 2024). Anterior to the parapophysis, there is a distinct fossa on the posterolateral surface of the prezygapophysis (Fig. 2A, 4A-B), similar to *Craterosaurus* (Galton 1981). In dorsal view, there is a midline ridge posterior to the prezygapophysis, connecting the base of the neural spine (Fig. 4C). The diapophyses extend dorsolaterally and form a high angle to the horizontal in anterior view, similar to other stegosaurs except *Gigantspinosaurus* with almost horizontal projected diapophyses (Hao et al. 2018). The diapophyses are dorsoventrally compressed. In the lateral view, the diapophysis has two distinct dorsoventrally ridges: paradiapophyseal lamina (PPDL) and posterior centrodiapophyseal lamina (PCDL) linked to the parapophysis. In posterior view, the outline of the postzygapophyses is triangular. The articular facets of the postzygapophyses are flat and confluent on the midline forming a V-shaped wedge (Fig. 4D). The dorsal part of the neural spine of most dorsal vertebrae is missing to varying degrees (Fig. 4B, C). Although the neural spine of the preserved dorsal vertebrae 5 and 6 looks complete, it may have been eroded. The neural spine is transversely compressed. It is short and sub-triangular in outline with a shorter anterior margin and longer posterior margin in lateral view (Fig. 4B). The apices of the neural spines do not expand transversely (Fig. 4C), different from *Huayangosaurus* and *Stegosaurus* (Maidment et al. 2006, 2015; Zhou 1984).

**Figure 4.**
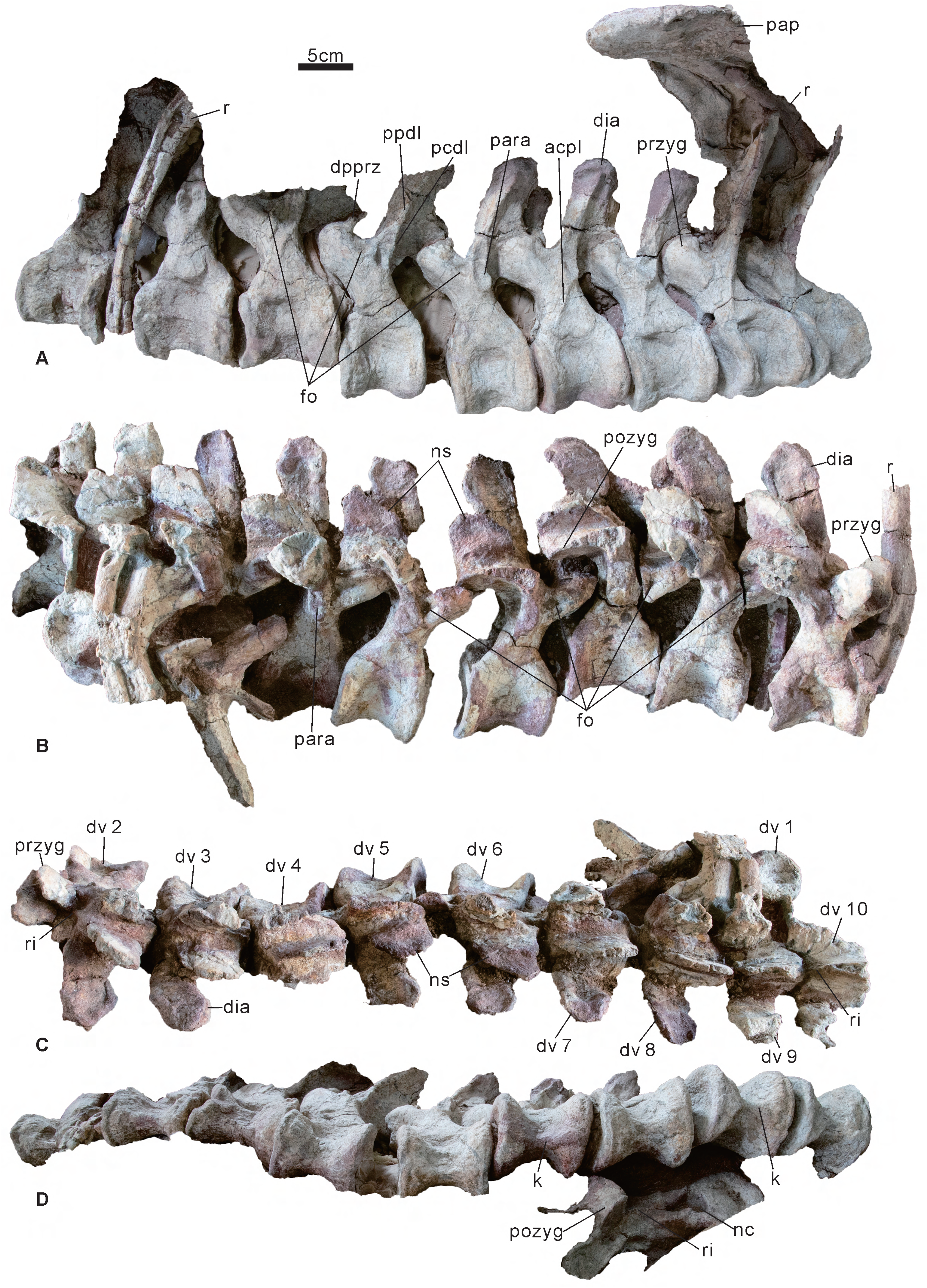
Ten dorsal vertebrae of *Angustungui qiketaiensis* (SS V 16001, holotype). A, left lateral view. B, right lateral view. C, dorsal view. D, ventral view. Abbreviations: acpl, anterior centroparapophyseal lamina; dia, diapophysis, dpprz, dorsal process of the prezygapophysis; dv, preserved dorsal vertebra; fo, fossa; k, keel; nc, neural canal; ns, neural spine; pap, preacetabular process; para, parapophysis; pcdl, posterior centrodiapophyseal lamina; pozyg, postzygapophysis; ppdl, paradiapophyseal lamina; przyg, prezygapophysis; r, rib; ri, ridge. Scale = 5 cm.

*Dorsal ribs*: Several dorsal ribs are preserved (Fig. 2). They are generally medially curved, with a reduced tuberculum, well-developed capitulum, and long shaft. The capitula and tubercula are compressed anteroposteriorly. Cross-sections of the proximal and distal halves of the shaft are L-shaped and oval. The last dorsal rib fused laterally to the medial margin of the preacetabular process of the ilium (Fig. 2A, 4A, 7B).

*Synsacrum and ribs*: The fused synsacrum consists of one dorsosacral vertebra and four sacral vertebrae, and they are fused to the ilia (Fig. 7B). The number of fused vertebrae forming the synsacrum varies among stegosaurs, even within the same genus such as *Dacentrurus*, *Kentrosaurus* and *Stegosaurus* (Galton 1982, 1985, 1991; Maidment et al. 2015; Sánchez-Fenollosa et al. 2024). The centre of the synsacrum is wider than it is long. The posterior articular facet of the last sacral vertebra is concave and has an oval outline with the long axis trending lateromedially. In the synsacrum series, the centre becomes progressively wider and the ventral keels are absent (Fig. 7B). The dorsosacral vertebra rib fused medially to the centrum, extended anterolaterally, and fused laterally to the medial margin of the preacetabular process of the ilium (Fig. 7B), similar to *Stegosaurus* (Maidment et al. 2015), unlike *Dacentrurus*, whose dorsosacral ribs fuse to dorsal margins of the first true sacral vertebra and *Yanbeilong* which fused to neither the first true sacral vertebra nor the ilium (Galton 1991; Jia et al. 2024). The sacral ribs fused medially to the centre, extend laterally and fused laterally to the medial surface of the ilium (Fig. 7B), unlike *Chungkingosaurus* and *Yanbeilong* have sacral ribs extend posterolaterally (Jia et al. 2024; Maidment and Wei 2006). The sacral ribs are anteroposteriorly expanded medially and laterally. The sacral vertebrae ribs form three large, oval fenestrae on each side in ventral view but the dorsal shield is solid without foramina (Fig. 7A, B), similar to *Dacentrurus*, *Kentrosaurus*, *Wuerhosaurus homheni*, *Stegosaurus*, *Hesperosaurus* and *Yanbeilong* (Carpenter et al. 2001; Dong 1990; Galton 1982, 1991; Jia et al. 2024; Maidment et al. 2015; Sánchez-Fenollosa et al. 2024), unlike the dorsal shield perforated by foramina between the sacral ribs in *Huayangosaurus*, *Gigantspinosaurus* and *Tuojiangosaurus* (Hao et al. 2018; Maidment et al. 2006; Maidment and Wei 2006; Zhou 1984). However, some fossae on the dorsal shield are symmetrically distributed on both sides of the neural spines (Fig. 7A). The neural spines of the dorsosacral vertebra and the first three sacral vertebrae are missing. Only the base of the neural spine of the last sacral vertebra is preserved, and it is hollow (Fig. 7A).

*Caudal vertebrae*: Seven nearly complete anterior caudal vertebrae are preserved, including two articulated caudal vertebrae (Fig. 5A), and five free caudal vertebrae (Fig. 5B-F). The two articulated anterior caudal vertebrae near the last sacral vertebra (Fig. 2), indicate they may be the first and second caudal vertebrae. The centrum is wider than long, and the anterior and posterior articular surfaces have a subcircular outline. The caudal centra are amphicoelous and the posterior articular surface is very deeply concave in the first and second caudal vertebra (Fig. 5Ab). The ventral surface of the centra is smooth, gently concave and lacking keel and chevron facets. The transverse processes of the first caudal vertebra are posterolaterally directed and anteroposteriorly compressed (Fig. 5Aa). The base of the transverse processes of the first caudal vertebra strongly expanded dorsoventrally and the dorsal margin has a lateromedial ridge in anterior and posterior views (Fig. 5A). Ventral to the ridge is a concave area, unlike *Dacentrurus, which* forms a proximodorsal canal of the transverse process at the same area (Sánchez-Fenollosa et al. 2024). In other caudal vertebrae, the transverse processes are strongly posteroventrally oriented and are not anteroposteriorly compressed, different from *Kentrosaurus* that the transverse processes of the anterior caudal vertebrae are laterally oriented (Galton 1982). The dorsal process of the transverse processes is present in the caudal vertebrae, and it is proximal to the centrum (Fig. 5Ba, b and Ea, b), similar to *Dacentrurus*, *Loricatosaurus*, *Stegosaurus* and *Hesperosaurus* (Carpenter et al. 2001; Galton 1985; Maidment et al. 2015; Sánchez-Fenollosa et al. 2024). The neural canal has a sub-circular outline in anterior and posterior views. The prezygapophyses project craniodorsally (Fig. 5Dc), unlike *Huayangosaurus*, which projects cranially (Zhou 1984). The articular facets of the prezygapophyses are flat, oval and face dorsomedially (Fig. 5De). The postzygapophyses extend caudally over the posterior facet of the centrum (Fig. 5Fd), unlike *Huayangosaurus*, *Kentrosaurus*, *Stegosaurus* and *Bashanosaurus*, in which the postzygapophyses do not extend caudally over the posterior facet of the centrum (Dai et al. 2022; Galton 1982; Maidment et al. 2015; Zhou 1984). The articular facets of the postzygapophyses are flat and face ventrolaterally (Fig. 5Fb). The neural spine extends posterodorsally and its height is greater than the height of the centrum (Fig. 5A-C), unlike *Huayangosaurus*, in which the neural spine height of the anterior caudal vertebrae is nearly equal to the height of the centrum (Zhou 1984). Among the seven caudal vertebrae, the neural spines of the first four project at a high angle to the horizontal in lateral view (Fig. 5A, Bc-d, Cc-d), but the neural spines of the last three project at a low angle (Fig. 5Dc-d, Ec-d, Fc-d). In anterior view, a dorsoventral midline ridge is present on the lower half of the neural spine (Fig. 5Ba, Ca, Ea), similar to *Miragaia* (Costa and Mateus 2019). The top of the neural spine is transversely expanded and bulbously swollen but does not bifurcate (Fig. 5Ca-b), similar to *Kentrosaurus*, *Hesperosaurus* and *Yanbeilong* (Carpenter et al. 2001; Galton 1982; Jia et al. 2024).

**Figure 5.**
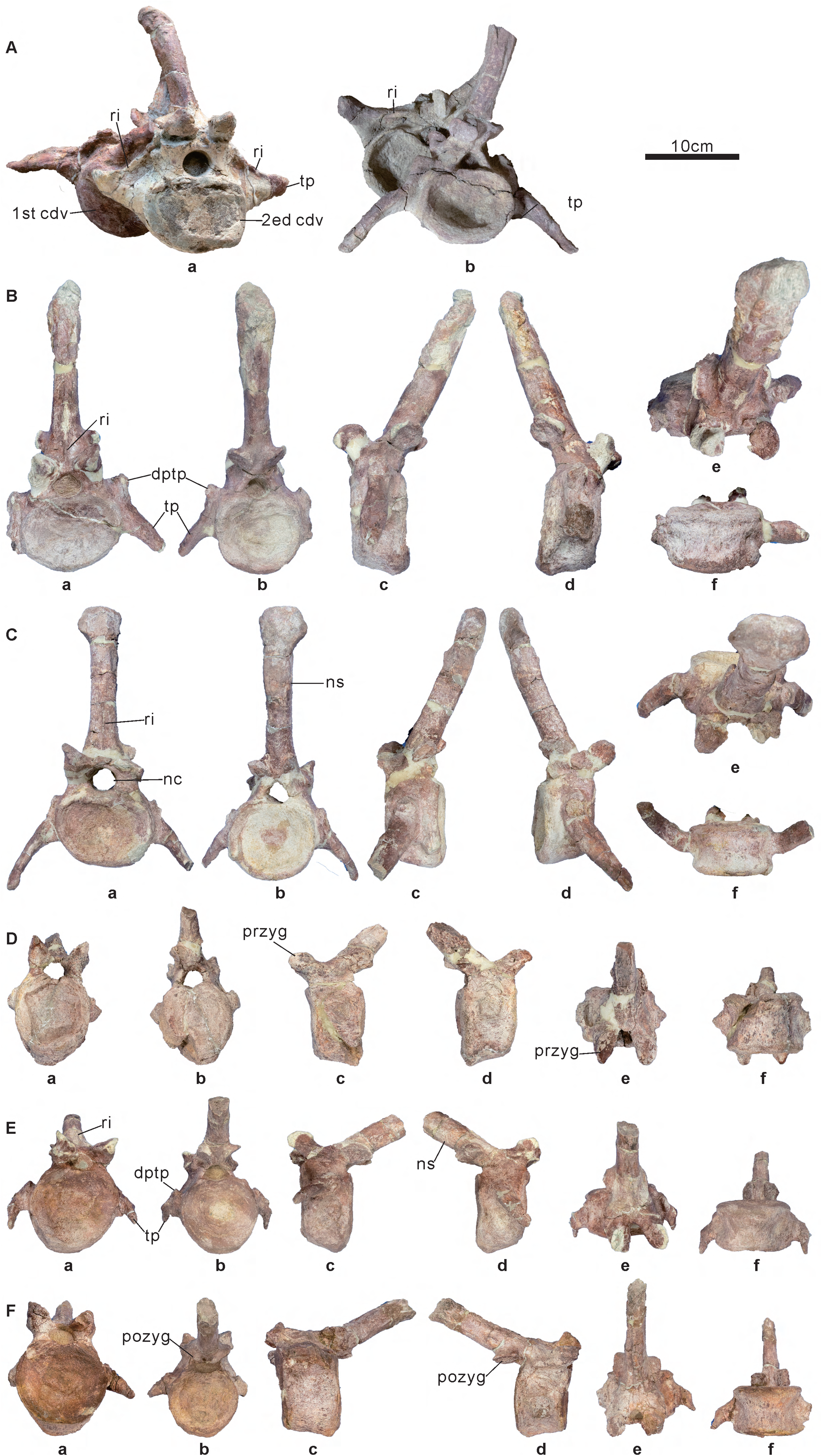
Seven anterior caudal vertebrae of *Angustungui qiketaiensis* (SS V 16001, holotype). A, first and second caudal vertebrae. B-F, preserved caudal vertebra 3 to 7. a, anterior view. b, posterior view. c, left lateral view. d, right lateral view. e, dorsal view. f, ventral view. Abbreviations: cdv, caudal vertebra; dptp, dorsal process of the transverse process; nc, neural canal; ns, neural spine; pozyg, postzygapophysis; przyg, prezygapophysis; ri, ridge; tp, transverse process. Scale = 10 cm.

#### Pectoral girdle

*Scapula*: The holotype (SS V16001) preserved a right scapula, missing the acromial region and most of the contact with the coracoid (Fig. 6A-D). The paratype (SS V16002) preserved a more complete right scapula, with only the dorsal part of the distal end of the blade and the dorsal part of the proximal plate missing (Fig. 6E-H). The paratype is slightly larger than the holotype. The proximal plate is gently concave in lateral view and flat in medial view (Fig. 6E-F). It forms the coracoid articulation anteriorly, the acromial process dorsally and the glenoid ventrally. The coracoid articulation is straight. The glenoid is concave and faces anteroventrally (Fig. 6D, F, H). In lateral view, the acromial process projects dorsally and its posterior margin forms an angle of approximately 90 degrees with the blade, similar to *Wuerhosaurus homheni* and *Stegosaurus* (Dong 1973; Maidment et al. 2015), but unlike *Huayangosaurus* and *Bashanosaurus,* the proximal plate is rounded posterodorsally (Dai et al. 2022; Maidment et al. 2006; Zhou 1984). The scapular blade is gently convex laterally and flat medially (Fig. 6A, B, E, F). The dorsal and ventral margins of the blade are sub-parallel along their entire lengths, similar to *Kentrosaurus* and *Stegosaurus* (Hennig 1915; Maidment et al. 2015), but different from *Bashanosaurus* that the blade is distally expanded (Dai et al. 2022). There is no evident sign of articulation with the parascapular spine.

**Figure 6.**
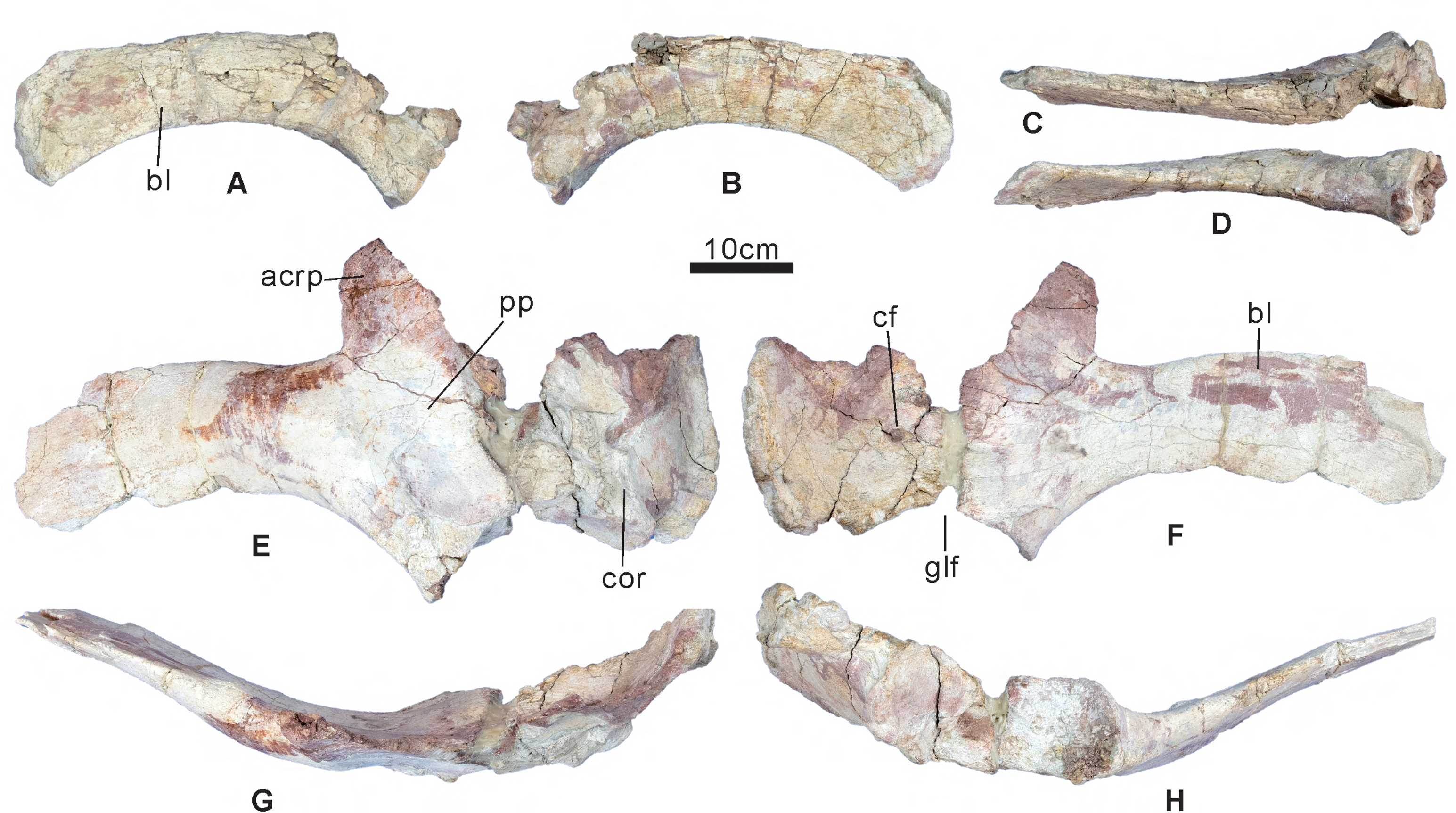
Pectoral girdle of *Angustungui qiketaiensis*. A-D, right scapula of the holotype (SS V16001). E-F, right scapula and coracoid of the paratype (SS V16002). A, E, lateral view. B, F, medial view. C, G, dorsal view. D, H, ventral view. Abbreviations: acrp, acromial process; bl, scapula blade; cf, coracoid foramen; cor, coracoid; glf, glenoid fossa; pp, scapula proximal plate. Scale = 10 cm.

*Coracoid*: The paratype (SS V16002) preserves a nearly complete right coracoid (Fig. 6E-H) not fused to the scapula. The fusion of coracoids and scapulae varied among *Stegosaurus* and *Kentrosaurus* and was likely ontogenetic (Maidment et al. 2015), but in the case of *Miragaia longicollum* type, the right scapulocoracoid is fused but the left is not (Mateus et al. 2009). In lateral view, the coracoid has a subcircular outline, similar to *Stegosaurus* and *Bashanosaurus* (Dai et al. 2022; Maidment et al. 2015), but different from *Huayangosaurus* and *Wuerhosaurus homheni* that the coracoid is longer anteroposteriorly than dorsoventrally high (Dong 1973; Maidment et al. 2006; Zhou 1984). The area of the proximal plate of the scapula is larger than the area of the coracoid, in contrast to *Huayangosaurus* and *Bashanosaurus* (Dai et al. 2022; Maidment et al. 2006; Zhou 1984). The coracoid foramen is oval in outline and is located in the posterior part of the coracoid at approximately mid-height (Fig. 6F).

#### Pelvic girdle

*Ilium*: The nearly complete ilia are preserved and fused to the synsacrum, but the preacetabular process of the right ilium has been deformed by compression (Fig. 7A, B). The preacetabular process has an inverted C-shaped cross-section that is laterally convex and medially concave, as in other stegosaurs. It projects roughly parallel to the parasagittal plane (Fig. 7A, B), similar to *Huayangosaurus* (Maidment et al. 2006; Zhou 1984), but different from *Dacentrurus*, *Stegosaurus* and *Yanbeilong* in that it diverges significantly from the parasagittal plane (Galton 1991; Jia et al. 2024; Maidment et al. 2015). In ventral view, the anterior tip of the preacetabular process is rounded and the ventrolateral margin of the preacetabular process is nearly straight (Fig. 7B). In lateral view, the preacetabular process lies approximately horizontally (Fig. 7C), similar to *Dacentrurus* and *Kentrosaurus* (Galton 1982, 1985, 1991). It differs from *Huayangosaurus*, *Wuerhosaurus ordosensis* and *Yanbeilong* in that the preacetabular process is strongly angled ventrally (Dong 1993; Jia et al. 2024; Maidment et al. 2006; Zhou 1984). The supra-acetabular process extends laterally and is hypertrophied to form a wing-like flange, similar to most stegosaurs. The supra-acetabular process and the preacetabular process form a gentle arc in dorsal and ventral views (Fig. 7A, B), unlike *Stegosaurus*, which forms a right angle (Maidment et al. 2015). The acetabulum lies ventromedial to the supra-acetabular process and is deeply concave. Its anterior margin is formed by an anteroventrally extended pubic peduncle and its posterior margin is formed by the ventrally extending ischiadic peduncle (Fig. 7C). There is a ventromedial flange between the pubic peduncle and ischiadic peduncle backing the acetabulum (Fig. 7C), which is absent in other stegosaurs except *Kentrosaurus* (Galton 1982). In dorsal view, the posterior iliac process is concave and its margin is convex (Fig. 7A). The posterior iliac processes have a tapered shape at the distal end, similar to *Huayangosaurus* and *Kentrosaurus* (Galton 1982; Maidment et al. 2006; Zhou 1984). In ventral view, an anteroposteriorly extended keel is present on the posterior iliac processes. There is a medial process medial to the keel, similar to *Hesperosaurus* and *Yanbeilong* (Carpenter et al. 2001; Jia et al. 2024).

**Figure 7.**
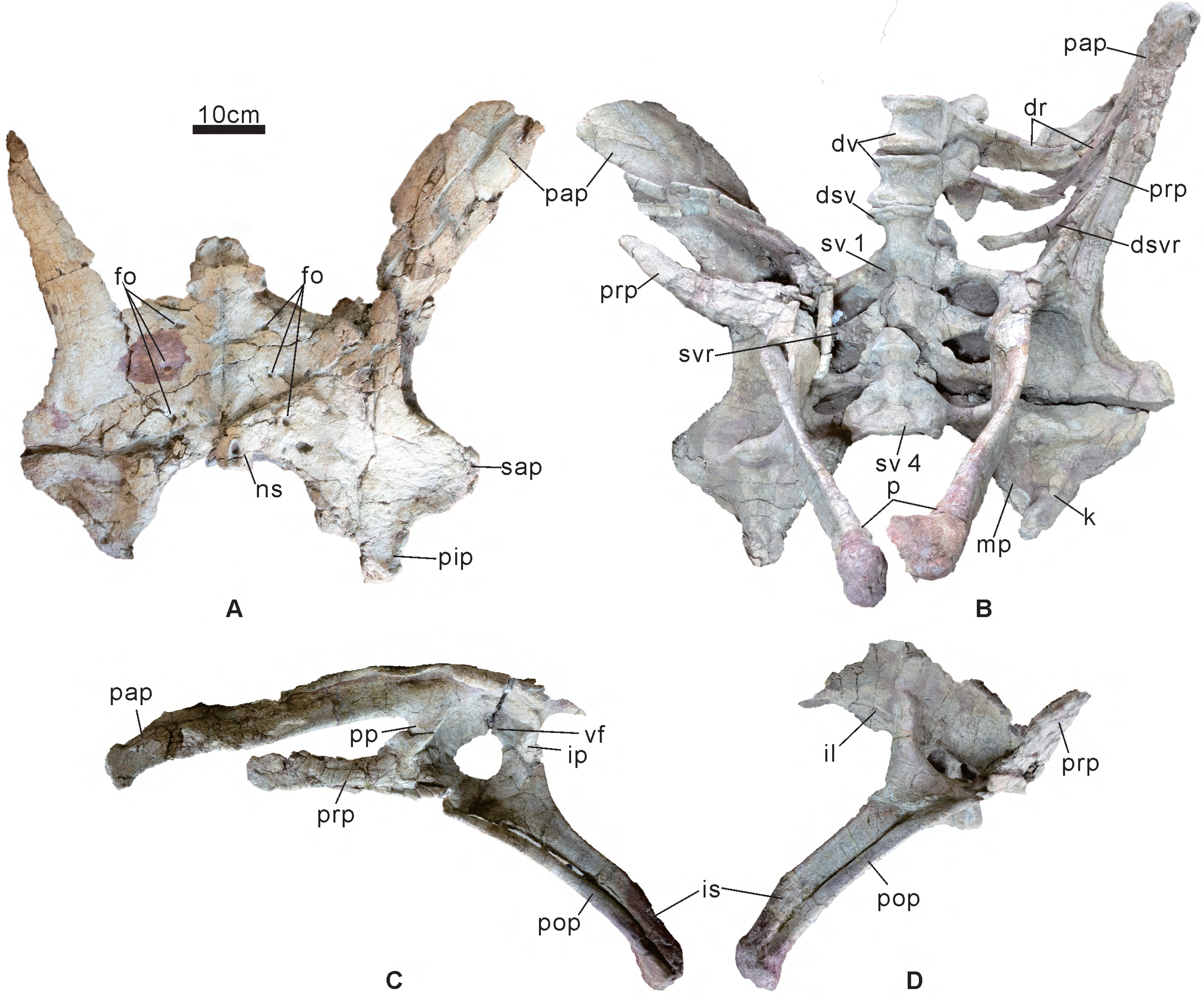
Synsacrum and pelvic girdle of *Angustungui qiketaiensis* (SS V16001). A, ilio-sacral block in dorsal view. B, synsacrum and pelvic girdle in ventral view. C, left pelvic girdle in lateral view. D, right pelvic girdle in lateral view. Abbreviations: dr, dorsal rib; dsv, dorsosacral vertebra; dsvr, dorsosacral vertebra rib; dv, dorsal vertebra; fo, fossa; ip, ischiadic peduncle; is, ischium; il, ilium; k, keel; mp, medial process; ns, neural spine; p, pubis; pap, preacetabular process; pip, posterior iliac process; pop, pubic shaft; pp, pubic peduncle; prp, prepubis; sap, supra-acetabular process; sv, sacral vertebrae; svr, sacral vertebrae rib; vf, ventromedial flange. Scale = 10 cm.

*Ischium*: Both ischia are well preserved (Fig. 7C, D). In the dorsal part, the pubic peduncle is situated anteriorly and the iliac peduncle is situated posteriorly. There is a deep, concave notch between the two peduncles that forms the part of the acetabular margin, similar to other stegosaurs. The dorsal surface of the shaft of the ischium has a distinct angle at approximately mid-length, similar to *Kentrosaurus* (Galton 1982), unlike *Dacentrurus* the shaft is straight (Galton 1991; Sánchez-Fenollosa et al. 2024). Ventral to the angle, the shaft of the ischium is tapered posteroventrally, but is transversely expanded (Fig. 7D). The distal end of the shaft is fused to the pubis.

*Pubis*: Both pubes are preserved. The left pubis is nearly complete (Fig. 7B, C), and the prepubis of the right pubis is deformed by compression (Fig. 7B, D). The prepubis extends anteriorly and is transversely compressed. The dorsal and ventral margins of the prepubis are sub-parallel along its entire length, forming a blunt anterior end in lateral view (Fig 71C), unlike *Dacentrurus* in which the anterior end of the prepubis is dorsally expanded (Galton 1985, 1991). The acetabular region is fused to the pubic peduncles of the ischium and the pubis (Fig 71C). It is transversely compressed and faces completely laterally, like other stegosaurs. It is difficult to determine the boundaries of the obturator notch because of the fractured nature of the bone. The pubic shaft extends posteroventrally and is slightly concave posterodorsally laterally (Fig 71C, D). The pubic shaft is much more slender than the prepubic and ischial shaft (Fig 71C, D). The distal end of the shaft is fused to the ischium and expanded transversely (Fig 71B, C, D).

#### Limb

*Femur*: The complete left femur is preserved (Fig. 8A-F). The femoral head extends dorsomedially and the greater trochanter lies lateral to the femoral head. In dorsal view, the femur is convex and sub-rectangular in outline (Fig. 8E). The femoral shaft is straight. The anterior and posterior surfaces of the shaft are depressed by crushing (Fig. 8A, B). The fourth trochanter is present on the posteromedial surface of the femur (Fig. 8B, C). It is a rugose ridge, similar to *Huayangosaurus* and *Bashanosaurus* (Dai et al. 2022; Zhou 1984). Two femoral epicondyles are located at the distal end of the femur. There is a deep intercondylar groove between the two femoral epicondyles in posterior view (Fig. 8B).

**Figure 8.**
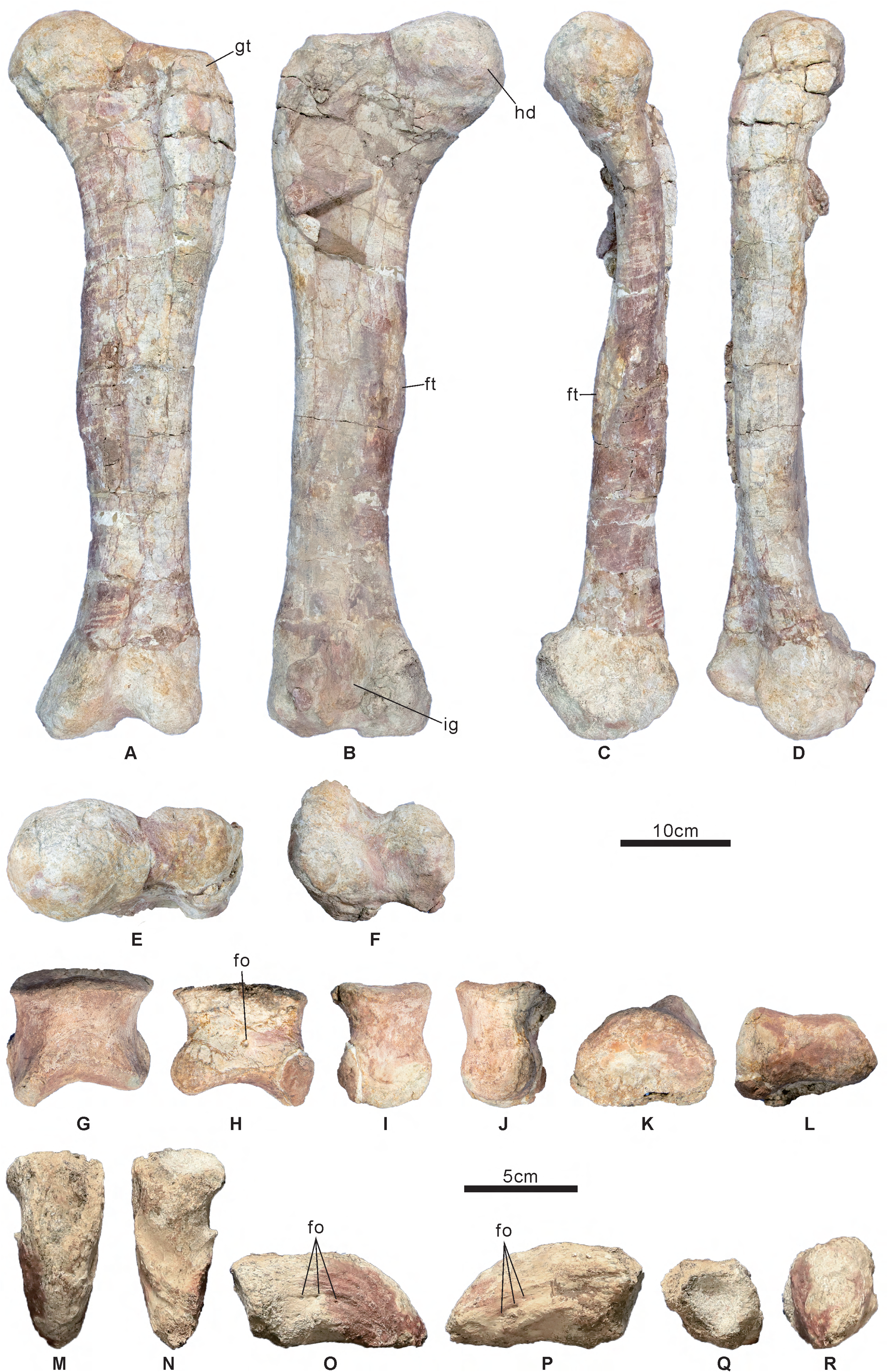
Limb of *Angustungui qiketaiensis* (SS V16001). A-F, left femur. G-L, non-terminal phalanx. M-R, ungual phalanx. A, G, R, anterior view. B, H, Q, posterior view. C, I, O, medial view. D, J, P, lateral view. E, K, M, dorsal view. F, L, N, ventral view. Abbreviations: fo, foramen; ft, fourth trochanter; gt, greater trochanter; hd, head; ig, intercondylar groove. Scale = 10 cm, (A–F), 5 cm (G-R).

*Non-terminal phalanx*: A complete non-terminal (non-ungual) phalanx is preserved (Fig. 8G-L). The original burial position of the phalanx and ungual is anterior to the left parascapular spine (Fig. 2), which suggests that the phalanx and ungual may belong to the left manus. In dorsal view, the proximal articular surface is D-shaped in outline with a straight posterior margin (Fig. 8K). In anterior and posterior views, its dorsal margin is straight and the lateral and medial margins are concave (Fig. 8G, H). The distal articular surface is saddle-shaped and has a sub-rectangular outline in ventral view (Fig. 8G, L). In posterior view, there is a foramen in the centre (Fig. 8H).

*Ungual phalanx*: One nearly complete ungual phalanx is preserved (Fig. 8M-R). The proximal articular surface is concave and sub-circular outline (Fig. 8Q). In dorsal and ventral views, the ungual tapers towards its distal tip and forms a blunt point at the end (Fig. 8M, N). In lateral and medial views, the ungual is slightly curved towards its tip, with a convex dorsal margin and a gently concave ventral margin (Fig. 8O, P). The lateral and medial surfaces display curved rows of foramina (Fig. 8O, P), similar to the thyreophoran *Scelidosaurus* (Norman 2020). The dorsoventral height and the mediolateral width of the ungual are nearly equal (Fig. 8R), which supported relatively narrow claw-shaped, unlike other stegosaurs with hoof-shaped ungual phalanges.

#### Dermal armor

*Parascapular spine*: The preserved complete spine is interpreted as a left parascapular spine (Fig. 9A-C). The parascapular spine is dorsoventrally compressed, and consists of a broad basal plate and a long spine (Fig. 9C). The basal plate is sub-triangular with a flat and bluntly posterior margin in dorsal and ventral views, on the same plane than the spine (Fig. 9A-B). In contrast, the basal plate is sub-quadrangular in *Loricatosaurus* and sub-circular in *Kentrosaurus* (Galton 1982, 1985) and sub-perpendicular to the orientation of the spine. In *Gigantspinosaurus*, the posterior margin of the basal plate is a more sharply pointed (Hao et al. 2018). The spine extends posterolaterally and tapers to the distal end. It has a gently curved anterolateral margin and a straight posteromedial margin. The dorsal surface of the spine is convex and the ventral surface is concave forming a deep groove (Fig. 9B). The base of the spine length is more than one-third of the lateral margin of the basal plate length (Fig. 9A-B), which is stronger than in *Gigantspinosaurus* (Hao et al. 2018). Proportionally, base of the spine is nearly the same length of the scapular blade.

**Figure 9.**
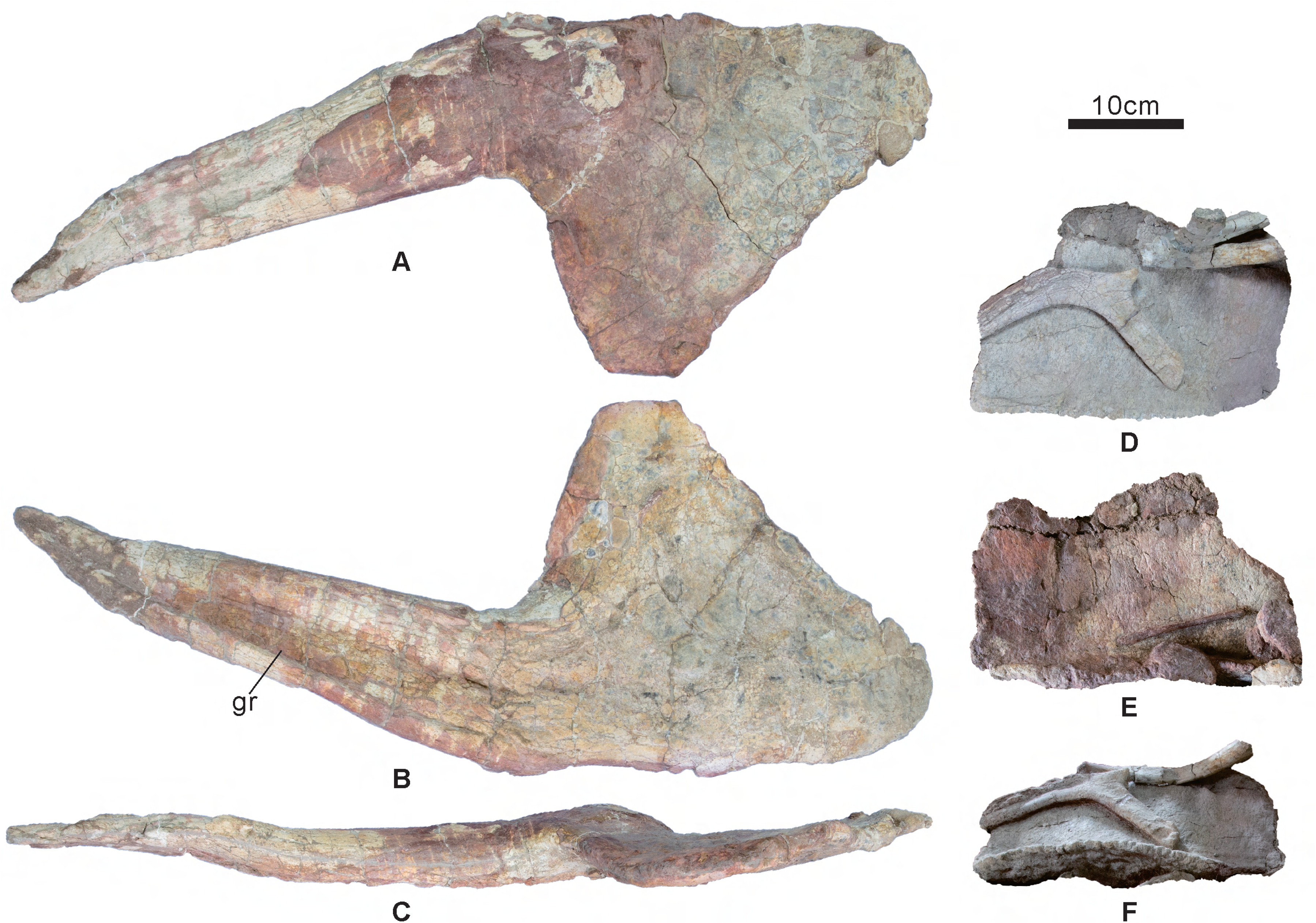
Dermal armors of *Angustungui qiketaiensis* (SS V16001). A-C, left parascapular spine. D-F, Dorsal plate. A, dorsal view. B, F, ventral view. C, D, left lateral view. E, right lateral view. Abbreviations: gr, groove. Scale = 10 cm.

*Plate*: One nearly complete dorsal plate is preserved (Fig. 2, 9D-F). The plate has a sub-rectangular outline in lateral view. The length of the plate is nearly four times the length of the dorsal centrum and half the length of the femur (Fig. 2). The ventral margin is convex and rugose, and transversely compressed dorsally ((Fig. 9F), similar to *Wuerhosaurus homheni* (Dong 1990), different from *Huayangosaurus*, *Kentrosaurus* the dorsal plates have a thick central portion like a modified spine (Galton 1982; Zhou 1984).

### Phylogenetic analysis

#### Revised character scores for Chinese stegosaurs

In the phylogenetic analysis of Raven and Maidment (2017), there were several clerical mistakes, that are revised here. First, in the character list, the definitions of Ch. 7 and Ch. 8 are the same. Ch. 8: Dorsal vertebrae, ‘centrum height to neural arch height ratio coded continuously’ changed to ‘centrum height to neural canal height ratio coded continuously’. Ch. 68: Anterior caudal vertebrae, dorsal process on transverse process proximal to centrum (0); ‘distal to centrum (2)’ changed to ‘distal to centrum (1)’. Ch. 73: Caudal vertebrae, ‘postzygapophyses extend cranially over caudal articular facet (0); do not (1)’ changed to ‘postzygapophyses extend caudally over caudal articular facet (0); do not (1)’. Then, the definition of Ch. 67: Anterior caudal vertebrae, dorsal process on transverse process absent (0); present (1). However, *Loricatosaurus* and *Alcovasaurus* are scored for 2 in the character-taxon matrix. The score is then changed to 1 for both *Loricatosaurus* and *Alcovasaurus*. Finally, Ch. 67 and Ch. 68 are two related morphological. If Ch. 67 scored 0 (the dorsal process on the transverse process of anterior caudal vertebrae is absent), Ch. 68 should score ?. However, *Scutellosaurus*, *Emausaurus*, *Scelidosaurus*, *Huayangosaurus*, *Gigantspinosaurus*, *Gastonia*, *Sauropelta* and *Yanbeilong* scored 0 for Ch. 67, but scored 0 for Ch. 68 in the character-taxon matrix. The scores of Ch. 68 of these taxa changed to ?.

*Huayangosaurus taibaii*: we revised 10 scorings for *Huayangosaurus* based on the holotype IVPP 6728 and referred specimen ZDM 7001. Ch. 1: snout, depth: depth to length ratio of maxilla coded as continuous. 0.8 changed to 0.44 (depth 57mm, length 131mm, Fig. 10A). Ch. 2: teeth, number coded as meristic. 21 changed to 27 (Fig. 10B). Ch. 6: dorsal vertebrae, neural arch to neural canal height ratio as continuous.changed to 1.41 (neural arch height 38mm, neural canal height 27mm, Fig. 10E). Ch. 7: dorsal vertebrae, centrum height to neural arch height ratio coded continuously.changed to 1.37 (centrum height 52mm, Fig. 10E). Ch. 8: dorsal vertebrae, centrum height to neural canal height ratio coded continuously. 1.6 changed to 1.93 (Fig. 10E). Ch. 11: scapula, proximal plate area to coracoid area ratio coded continuously. 1.25 changed to 0.82 (proximal plate area 119 cm^2^, coracoid area 145 cm^2^, Fig. 10F). The proximal plate of the scapula is significantly smaller in area than the coracoid. Ch. 31: lacrimal, contacts prefrontal (0); doesn’t contact prefrontal (1). 1 changed to 0 (Fig. 10A). Ch. 56: axis, ventral margin in lateral view flat (0); concave (1).changed to 1 (Fig. 10C). Ch. 57: cervical vertebra 3, centrum ventral margin straight (0); concave upwards (1).changed to 0 (Fig. 10D). Ch. 68: anterior caudal vertebrae, dorsal process on transverse process proximal to centrum (0); distal to centrum (1). 0 changed to(dorsal process on transverse process is absent, Ch. 67 scores for 0).

**Figure 10.**
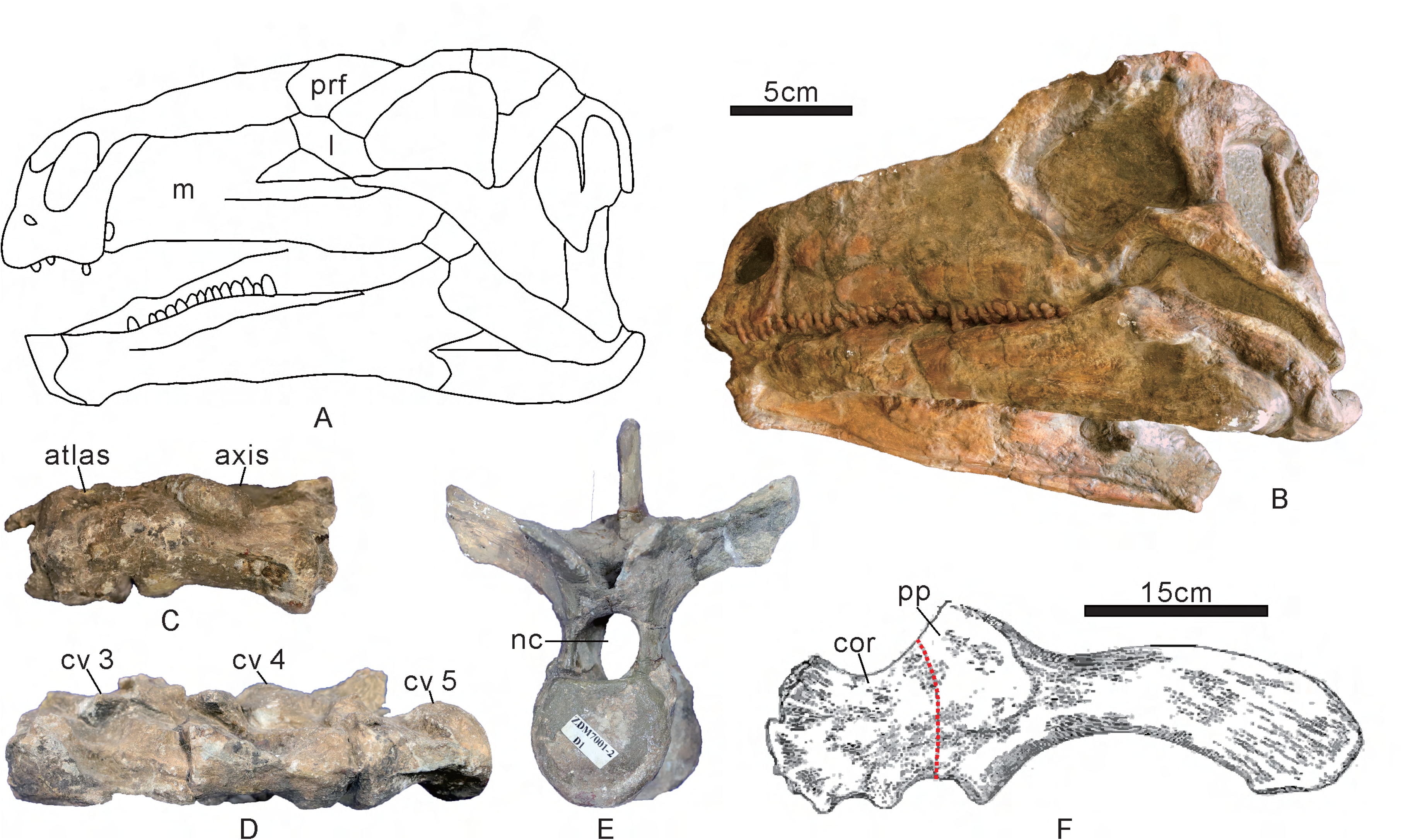
*Huayangosaurus taibaii*. A, sketch of the skull (ZDM 7001) in left lateral view, modified from Sereno and Dong (1992). B, skull (IVPP 6728) in left lateral view. C, atlas and axis (ZDM 7001) in left lateral view. D, cervical vertebra 3 to 5 (ZDM 7001) in left lateral view. E, dorsal vertebra 1 (ZDM 7001) in anterior view. F, left scapulocoracoid (ZDM 7001) in lateral view, modified from Maidment et al. (2006). Abbreviations: cor, coracoid; cv, cervical vertebra; l, lacrimal; m, maxilla; nc, neural canal; pp, proximal plate; prf, prefrontal. Scale = 5 cm (A–E), 15 cm (F).

*Tuojiangosaurus multispinus*: All the holotype specimens (CQMNH-CV 209) except the skull were mounted and displayed in the lobby of the Chongqing Natural History Museum. However, most bones have been skimmed with plaster so their original shape and anatomical details are difficult to determine. we revised 26 scorings for *Tuojiangosaurus* based on the holotype. Ch. 3: teeth, number of denticles on the mesial side of maxillary teeth.changed to 7 (Fig. 11A). Ch. 6: dorsal vertebrae, neural arch to neural canal height ratio as continuous.changed to 1.8 (neural arch height 54mm, neural canal height 30mm, Fig. 11C). Ch. 7: dorsal vertebrae, centrum height to neural arch height ratio coded continuously.changed to 0.98 (centrum height 53mm, Fig. 11C). Ch. 8: dorsal vertebrae, centrum height to neural canal height ratio coded continuously.changed to 1.77 (Fig. 11C). Ch. 12: humerus, ratio of width of distal end to minimum shaft width coded continuously.changed to 2.22 (distal end width 195mm, minimum shaft width 88mm, Fig. 11G). The left humerus is well preserved. Ch. 13: humerus, ratio of transverse width of distal end to length coded continuously.changed to 0.34 (length 575mm, Fig. 11G). Ch. 14: humerus, anterior iliac process length to humerus length coded continuously.changed to 0.9 (anterior iliac process length 515mm, Fig. 11G, J). The ilium is well preserved. Ch. 23: femur, length to humerus length ratio coded continuously.changed to 1.49 (femur length 855mm, Fig. 11G, K). The right femur is well preserved. Ch. 24: femur, length to tibia length ratio continuously.changed to 1.48 (tibia length 578mm, Fig. 11I, K). The right tibia is well preserved. Ch. 58: cervical vertebrae, longer anterposteriorly than wide transversely (0); wider than long (1).changed to 0 (Fig. 11B). The cervical vertebra only preserved centrum. Ch. 60: anterior dorsal vertebrae, prezygapophyses are separated and face each other dorsally (0); joined ventrally and face dorsomedially (1).changed to 1 (Fig. 11C). Ch. 61: dorsal vertebrae, cranial and caudal articular facets on centra flat to slightly concave (0); strongly convex (1).changed to 0 (Fig. 11C). Ch. 62: dorsal vertebrae, all centra longer than wide (0); wider than long (1).changed to 0 (Fig. 11C, D). Ch. 63: dorsal vertebrae, transverse processes project approximately horizontally (0); at a high angle to the horizontal (1).changed to 1 (Fig. 11C). Ch. 67: anterior caudal vertebrae, dorsal process on transverse process absent (0); present (1).changed to 0 (Fig. 11F). The caudal vertebra 3 is nearly well preserved, only the prezygapophyses is absent. Ch. 69: anterior caudal vertebrae, transverse processes on cd3 posteriorly are directed laterally (0); directed strongly ventrally (1).changed to 1 (Fig. 11F). Ch. 70: anterior caudal vertebrae, neural spine height less than or equal to the height of the centrum (0); greater than the height of the centrum (1).changed to 1 (Fig. 11F). Ch. 71: anterior caudal vertebrae, bulbous swelling at tops of neural spines absent (0); present (1).changed to 0 (Fig. 11F). Ch. 72: caudal vertebrae, prezygapophyses extend craniodorsally (0); extend cranially (1).changed to 1 (Fig. 11E). The caudal vertebrae 20 to 22 nearly are well preserved, only the neural spines are absent. Ch. 73: caudal vertebrae, postzygapophyses extend caudally over caudal articular facet (0); do not (1).changed to 1 (Fig. 11E). Ch. 74: caudal vertebrae, transverse processes on distal half of tail present (0); absent (1).changed to 0 (Fig. 11E). Ch. 75: caudal vertebrae, neural spines bifurcated (0); not bifurcated (1).changed to 1 (Fig. 11F). Ch. 76: posterior caudal vertebrae, centra are elongate (0); equidimensional (1).changed to 0 (Fig. 11E). Ch. 79: scapula, blade, distally expanded (0); parallel sided (1).changed to 0 (Fig. 11H). The blade of the left scapula is preserved. Ch. 82: humerus, triceps tubercle and descending ridge posterolateral to the deltopectoral crest absent (0); present (1).changed to 1 (Fig. 11F). Ch. 115: dorsal plates, have a thick central portion like a modified spine (0); have a generally transversely thin structure, except at the base (1).changed to 0 (Fig. 11L).

**Figure 11.**
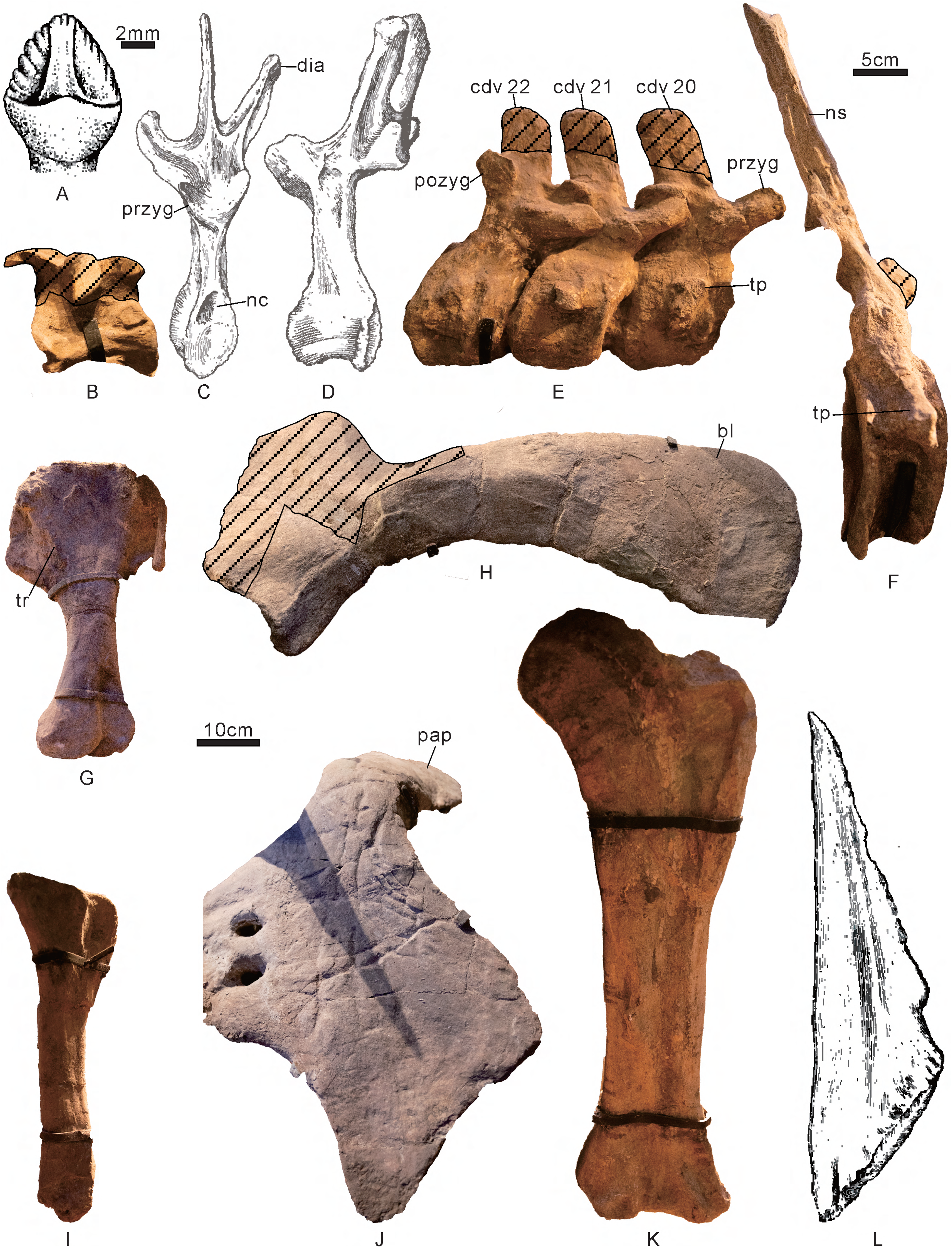
*Tuojiangosaurus multispinus* (CQMNH-CV 209). A, maxillary tooth in labial views, from Galton et al. (2004). B, posterior cervical vertebra in right lateral view. C-D, dorsal vertebra in anterior and left lateral views, from Dong et al. (1977). E, caudal vertebrae 20 to 22 in right lateral view. F, caudal vertebra 3 in right lateral view. G, left humerus in posterior view. H, left scapula in lateral view. I, right tibia in posterior view. J, right ilium in dorsal view. K, right femur in posterior view. L, dorsal plate in lateral view, from Dong et al. (1977). Abbreviations: bl, scapula blade; cdv, caudal vertebra; dia, diapophysis; nc, neural canal; ns, neural spine; pap, preacetabular process; pozyg, postzygapophysis; przyg, prezygapophysis; tp, transverse process; tr, triceps ridge. Striped regions indicate reconstruction. Scale = 2 mm (A), 5cm (B-F), 10cm (G-L).

*Chungkingosaurus jiangbeiensis*: All the holotype specimens (CQMNH-CV 206) except the skull were also mounted and displayed in the lobby of the Chongqing Natural History Museum and most bones have been skimmed with plaster. we revised 10 scorings for *Chungkingosaurus* based on the holotype. Ch. 24: femur, length to tibia length ratio continuously.changed to 1.17 (femur length 413mm, tibia length 354mm, Fig. 12E, F). Ch. 61: dorsal vertebrae, cranial and caudal articular facets on centra flat to slightly concave (0); strongly convex (1). 1 changed to 0 (Fig. 12A). Ch. 66: sacral rod vertebrae, keel present (0); absent (1).changed to 1 (Fig. 12B). Ch. 67: anterior caudal vertebrae, dorsal process on transverse process absent (0); present (1).changed to 0 (Fig. 12C). Ch. 69: anterior caudal vertebrae, transverse processes on cd3 posteriorly are directed laterally (0); directed strongly ventrally (1).changed to 0 (Fig. 12C). Ch. 72: caudal vertebrae, prezygapophyses extend craniodorsally (0); extend cranially (1).changed to 1 (Fig. 12C). Ch. 73: caudal vertebrae, postzygapophyses extend caudally over caudal articular facet (0); do not (1).changed to 0 (Fig. 12C). Ch. 74: caudal vertebrae, transverse processes on distal half of tail present (0); absent (1).changed to 1 (Fig. 12D). Ch. 104: femur, fourth trochanter prominent and pendant (0); present as a rugose ridge (1); absent (2).changed to 2 (Fig. 12E). Ch. 105: femur, anterior trochanter fusion to greater trochanter in adults - unfused (0); fused (1).changed to 0 (Fig. 12E). Ch. 104 and Ch. 105 are also described by Dong et al. 1983 (1983).

**Figure 12.**
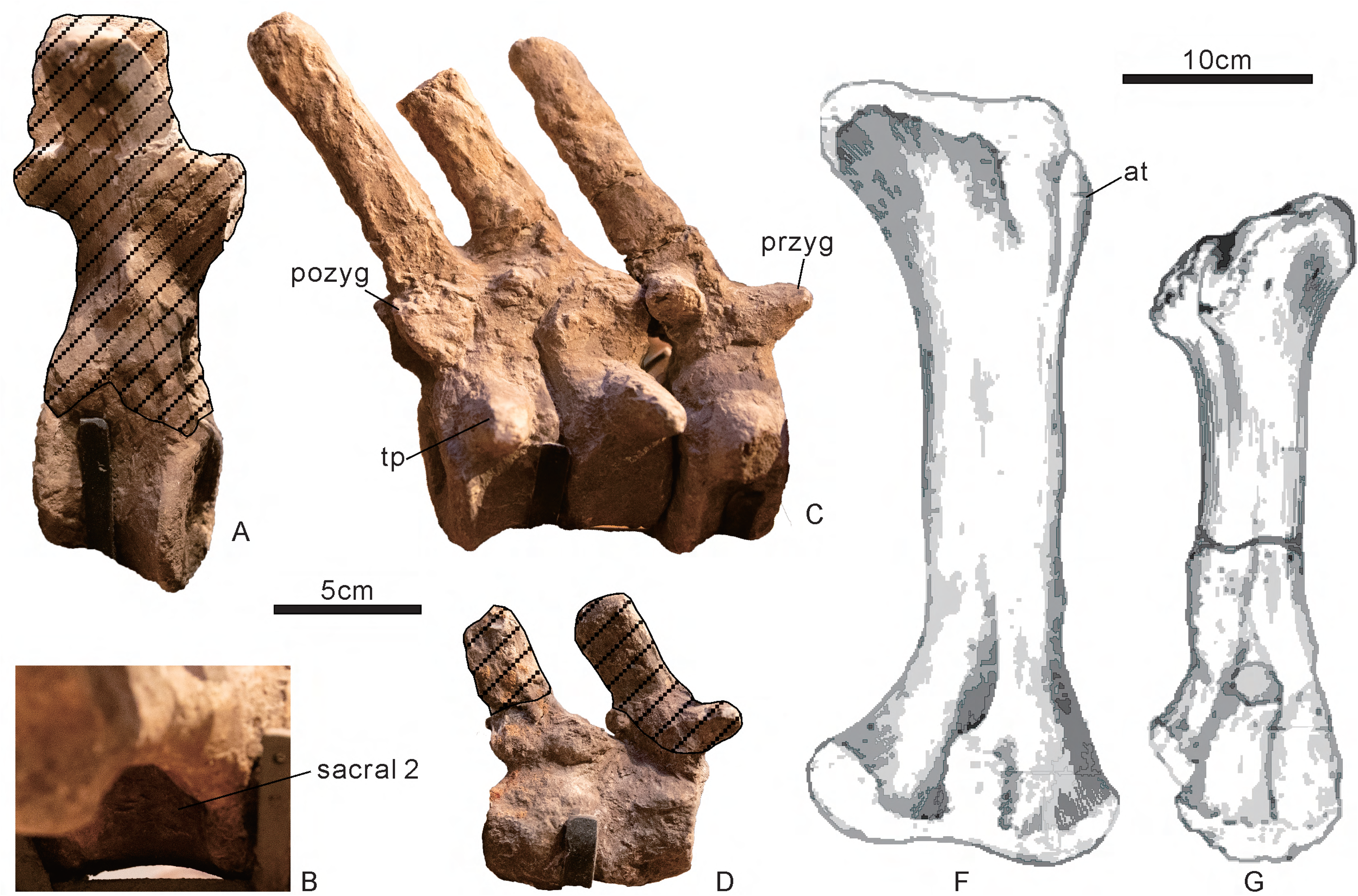
*Chungkingosaurus jiangbeiensis* (CQMNH-CV 206). A, middle dorsal vertebra in right lateral view. B, sacral vertebra 2 in right lateral view. C, three associated anterior caudal vertebrae in right lateral view. D, two associated posterior caudal vertebrae in right lateral view. E, right femur in posterior view. F, left tibia in lateral view. E and F from Dong et al. (1983). Abbreviations: at, anterior trochanter; pozyg, postzygapophysis; przyg, prezygapophysis; tp, transverse process. Striped regions indicate reconstruction. Scale = 5cm (A-D), 10cm (E-F).

*Gigantspinosaurus sichuanensis*: we revised 30 scorings for *Gigantspinosaurus* based on the holotype ZDM 0019. Ch. 5: cervical vertebrae, number coded as meristic.changed to 8 (Fig. 13C, D). Ch. 6: dorsal vertebrae, neural arch to neural canal height ratio as continuous.changed to 1.59 (neural arch height 38mm, neural canal height 27mm, Fig. 13I). Ch. 7: dorsal vertebrae, centrum height to neural arch height ratio coded continuously.changed to 1.12 (centrum height 52mm, Fig. 13I). Ch. 8: dorsal vertebrae, centrum height to neural canal height ratio coded continuously.changed to 1.78 (Fig. 13I). Ch. 11: scapula, proximal plate area to coracoid area ratio coded continuously.changed to 0.71 (the proximal plate of the scapula is complete with area 279cm^2^, coracoid area 395cm^2^, Fig. 13E). Ch. 12: humerus, ratio of width of distal end to minimum shaft width coded continuously.changed to 2.26 (distal end width 158mm, minimum shaft width 70mm, Fig. 13J). Ch. 14: humerus, anterior iliac process length to humerus length coded continuously.changed to 0.97 (anterior iliac process length 410mm, humerus length 424mm, Fig. 13F, J). Ch. 15: ulna, proximal width to length ratio coded continuously.changed to 0.41 (proximal width 153mm, length 372mm, Fig. 13J). Ch. 18: metacarpal II to humerus length ratio coded continuously.changed to 0.17 (metacarpal II length 74mm, humerus length 424mm, Fig. 13J). Ch. 19: ilium, anterior iliac process to acetabular length ratio coded continuously.changed to 2.1 (anterior iliac process length 410mm, acetabular length 195mm, Fig. 13F). Ch. 20: ilium, ratio of acetabular length to dorsoventral height of pubic peduncle of ilium coded continuously.changed to 3 (acetabular length 195mm, height of pubic peduncle of ilium 65mm, Fig. 13F). Ch. 44: dentary, postdentary bones greater in rostrocaudal length than dentary (0); shorter (1).changed to 1 (Fig. 13A). Ch. 48: tooth crowns, striations not confluent with denticles (0); confluent with denticles (1).changed to 0 (Fig. 13A). Ch. 49: tooth crowns, asymmetric (0); symmetric (1).changed to 0 (Fig. 13A). Ch. 56: axis, ventral margin in lateral view flat (0); concave (1).changed to 1 (Fig. 13B, C). Ch. 57: cv3, centrum ventral margin straight (0); concave upwards (1).changed to 1 (Fig. 13B). Ch. 58: cervical vertebrae, longer anterposteriorly than wide transversely (0); wider than long (1).changed to 0 (Fig. 13D). Ch. 59: posterior cervical vertebrae, postzygapophyses not greatly elongated (0); greatly elongated and project over the back of the posterior centrum facet (1).changed to 1 (Fig. 13C). Ch. 61: dorsal vertebrae, cranial and caudal articular facets on centra flat to slightly concave (0); strongly convex (1).changed to 0 (Fig. 13I). Ch. 68: anterior caudal vertebrae, dorsal process on transverse process proximal to centrum (0); distal to centrum (1). 0 changed to(dorsal process on transverse process is absent, Ch. 67 scores for 0). Ch. 72: caudal vertebrae, prezygapophyses extend craniodorsally (0); extend cranially (1).changed to 0 (Fig. 13H). Ch. 75. caudal vertebrae: neural spines bifurcated (0); not bifurcated (1).changed to 1 (Fig. 13G). Ch. 80: coracoid, sub-circular outline (0); anteroposteriorly longer than dorsoventrally high (1).changed to 1 (Fig. 13E). Ch. 83: radius, expanded transversely at proximal end (0); not expanded (1).changed to 0 (Fig. 13J). Ch. 84: metacarpals I and V, shorter than metacarpals II, III and IV (0); longer (1).changed to 0 (Fig. 13J). Ch. 85: ungual phalanges, manual and pedal unguals claw–shaped (0); hoof–shaped (1).changed to 1 (Fig. 13J). Ch. 93: Ilium, ventromedial flange backing the acetabulum absent (0); present (1).changed to 0 (Fig. 13F). Ch. 98: ischium, convex proximal margin within the acetabulum absent (0); present (1).changed to 0 (Fig. 13F). Ch. 99: ischium, dorsal surface of shaft is straight (0); has a distinct angle at approximately midlength (1).changed to 1 (Fig. 13F). Ch. 102: pubis, acetabular portion faces laterally, posteriorly and dorsally (0); faces wholly laterally (1).changed to 1 (Fig. 13F).

**Figure 13.**
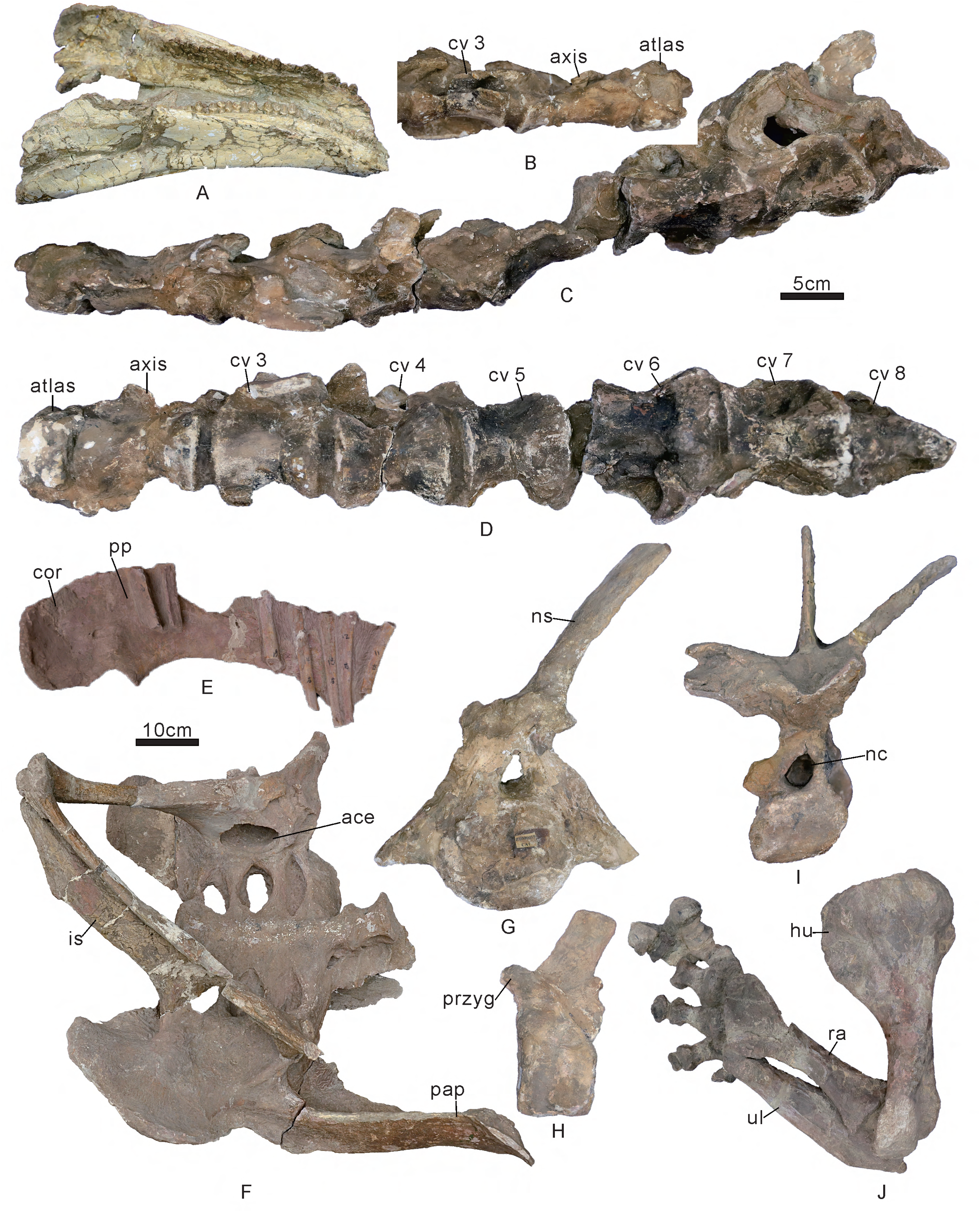
*Gigantspinosaurus sichuanensis* (ZDM 0019). A, mandible in dorsolateral view. B, atlas, axis and cervical vertebra 3 in right lateral view. C, cervical vertebra 1 to 8 in left lateral view. D, cervical vertebra 1 to 8 in ventral view. E, right scapulocoracoid in medial view. F, ilio-sacral block in ventrolateral view. G, caudal vertebra 1 in anterior view. H, caudal vertebra 18 in left lateral view. I, dorsal vertebra 12 in anterior view. J, forelimb in posterior view. Abbreviations: ace, acetabulum; cor, coracoid; cv, cervical vertebra; hu, humerus; is, ischium; nc, neural canal; ns, neural spine; pap, preacetabular process; pp, proximal plate; przyg, prezygapophysis; ra, radius; ul, ulna. Scale = 5 cm (A–D, G-I), 10 cm (E, F, J).

Other taxa. *Wuerhosaurus homheni*: we revised 11 scorings for *Wuerhosaurus homheni* based on the holotype IVPP 4006. Ch. 11: scapula, proximal plate area to coracoid area ratio coded continuously.changed to 1.31 (proximal plate area 739 cm^2^, coracoid area 565 cm^2^, Fig. 14D). Ch. 61: dorsal vertebrae, cranial and caudal articular facets on centra flat to slightly concave (0); strongly convex (1).changed to 0 (Fig. 14A). Ch. 77: scapula, acromial process in lateral view, convex upwards dorsally (0); quadrilateral with a posterordorsal corner (1).changed to 1 (Fig. 14D). Ch. 78: scapula, acromial process projects dorsally (0); projects laterally (1).changed to 0 (Fig. 14D). Ch. 79: scapula, blade, distally expanded (0); parallel sided (1).changed to 1 (Fig. 14D). Ch. 80: coracoid, sub-circular outline (0); anteroposteriorly longer than dorsoventrally high (1).changed to 1 (Fig. 14D). Ch. 81: coracoid, in lateral view, foramen present (0); notch present (1).changed to 0 (Fig. 14D). Ch. 91: ilium, posterior iliac process, distal shape tapers (0); blunt (1). 1 changed to 0 (Fig. 14E). Ch. 101: pubis, obturator notch is backed by posterior pubic process absent (0); present (1).changed to 0 (Fig. 14C). Ch. 102: pubis, acetabular portion faces laterally, posteriorly and dorsally (0); faces wholly laterally (1).changed to 1 (Fig. 14C). Ch. 103: pubis, anterior end of prepubis expanded dorsally absent (0); present (1).changed to 0 (Fig. 14C). *Jiangjunosaurus junggarensis*: we revised 5 scorings for *Jiangjunosaurus* based on the holotype IVPP V14724. Ch. 58: cervical vertebrae, longer anterposteriorly than wide transversely (0); wider than long (1).changed to 0 (Fig. 14F, G). Ch. 59: posterior cervical vertebrae, postzygapophyses not greatly elongated (0); greatly elongated and project over the back of the posterior centrum facet (1).changed to 1 (Fig. 14F). Ch. 61: dorsal vertebrae, cranial and caudal articular facets on centra flat to slightly concave (0); strongly convex (1). 0 changed to ?. The dorsal vertebrae are not preserved. Ch. 79: scapula, blade, distally expanded (0); parallel sided (1). 0 changed to ?. The blade of the scapula is not preserved (Fig. 14H). Ch. 81: coracoid, in lateral view, foramen present (0); notch present (1).changed to 1 (Fig. 14H). *Yanbeilong ultimus*: we revised 1 scoring for *Yanbeilong* based on the holotype SXMG V 00006. Ch. 68: anterior caudal vertebrae, dorsal process on transverse process proximal to centrum (0); distal to centrum (1). 0 changed to(dorsal process on transverse process is absent, Ch. 67 scores for 0).

**Figure 14.**
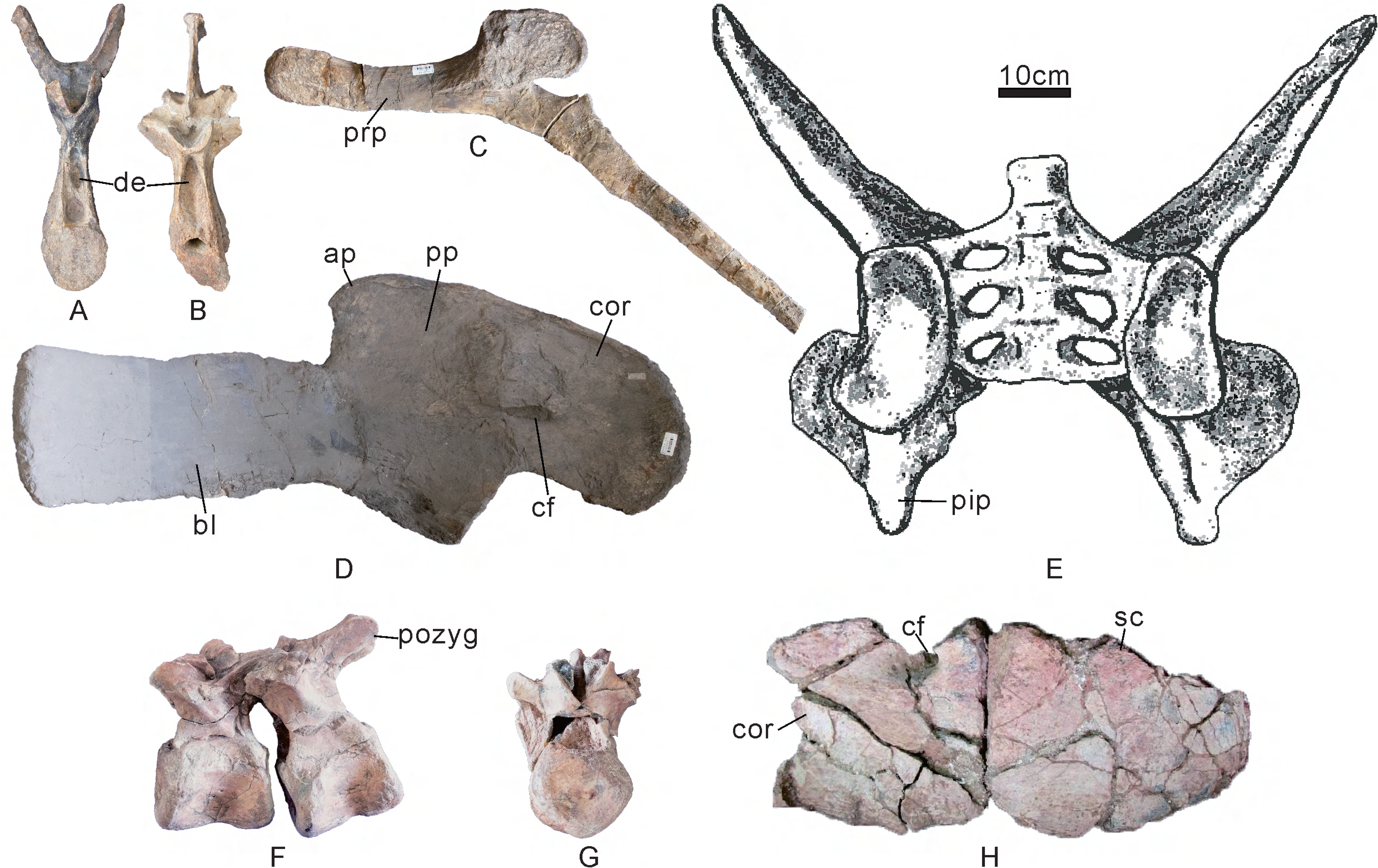
*Wuerhosaurus*. A, C, D, E, *Wuerhosaurus homheni* (IVPP 4006). B, *Wuerhosaurus ordosensis* (IVPP 6879). F-H, *Jiangjunosaurus junggarensis* (IVPP V14724). A, B, dorsal vertebra in anterior view. C, left pubis in lateral view. D, left scapulocoracoid in medial view. E, ilio-sacral block in ventral view, from Dong (1990). F, G, cervical vertebrae 7 and 8 in lateral view and anterior view. H, left scapulocoracoid in lateral view. Abbreviations: ap, acromial process; bl, scapula blade; cf, coracoid foramen; cor, coracoid; de, deep excavation; pip, posterior iliac process; pozyg, postzygapophysis; pp, proximal plate; prp, prepubis; sc, scapula. Scale = 10 cm.

Besides, Li et al. (2024a) added two new characters to the dataset. Ch. 9: dorsal vertebrae, neural spines length (measured at the base) to centrum length ratio coded continuously. Ch. 64: dorsal vertebrae, parapophyses are well developed that held on stalks at the base of the diapophyses; (0); poorly developed (1). We scored the two characters for *Yuxisaurus*, *Yanbeilong* and *Thyreosaurus*. *Yuxisaurus* (Yao et al. 2022): Ch. 9 scores(the neural spines of dorsal vertebrae are absent). Ch. 64 scores 0. *Yanbeilong* (Jia et al. 2024): Ch. 9 scores 0.48 (neural spines length 45mm, centrum length 94mm). Ch. 64 scores 1. *Thyreosaurus* (Zafaty et al. 2024): Ch. 9 scores 0.66 (neural spines length 53mm, centrum length 80mm). Ch. 64 scores 1.

#### Analysis results

The seven most parsimonious trees (MPTs) were recovered with tree lengths of 306.35 steps, a Consistency Index of 0.534 and a Retention Index of 0.634. In these MPTs, the clade of the Stegosauria is stable, including Huayangosauridae (Dong et al. 1982) and Stegosauridae (Marsh 1880). The Huayangosauridae consist of *Huayangosaurus*, *Bashanosaurus*, *Chungkingosaurus*, *Isaberrysaura*, *Gigantspinosaurus*, *Baiyinosaurus* and *Tuojiangosaurus*, which is supported by the five synapomorphies: ch. 2 (more than 27) the number of teeth; ch. 11 (less than 0.82) the ratio of proximal plate of scapula area to coracoid area; ch. 49 (0) the tooth crowns are asymmetric (0); ch. 51 (0) the premaxillary teeth are present and ch. 56 (1) the ventral margin of axis in lateral view is concave. Stegosauridae includes Stegosaurinae (Marsh 1880) and Dacentrurinae (Mateus et al. 2009) with *Miragaia* and *Dacentrurus* being well differentiated. The Stegosauridae consist of *Alcovasaurus*, *Kentrosaurus*, Dacentrurinae including *Adratiklit*, *Miragaia*, *Thyreosaurus* and *Dacentrurus,* and Stegosaurinae with *Jiangjunosaurus*, *Hesperosaurus*, *Angustungui*, *Loricatosaurus*, *Yanbeilong*, *Stegosaurus stenops*, and *Wuerhosaurus homheni*. The clade Stegosauridae is supported by the two synapomorphies: ch. 67 (1) the dorsal process on transverse process of the anterior caudal vertebrae is present and ch. 71 (1) the bulbous swelling at tops of neural spines of the anterior caudal vertebrae is present. *Angustungui* is recovered as the sister taxon of *Loricatosaurus*, which is supported by the two synapomorphies: ch. 69 (1) the transverse processes on cd3 posteriorly are directed strongly ventrally and ch. 114 (1) the parascapular spine is present. The seven MPTs differ only in their placement of *Yuxisaurus*, *Paranthodon* and the Ankylosaurs *Gastonia*, *Sauropelta* and *Euoplocephalus*. A strict consensus of the MPTs and the measures of support for clades are shown in Fig. 15. The strict consensus of the *Yuxisaurus*, *Paranthodon* and the Ankylosaurs are poorly resolved, which due to labile phylogenetic position of *Yuxisaurus* and *Paranthodon* (Raven and Maidment 2018; Yao et al. 2022). The revised character list and data matrix can be found in supplementary 2 and 3.

**Figure 15.**
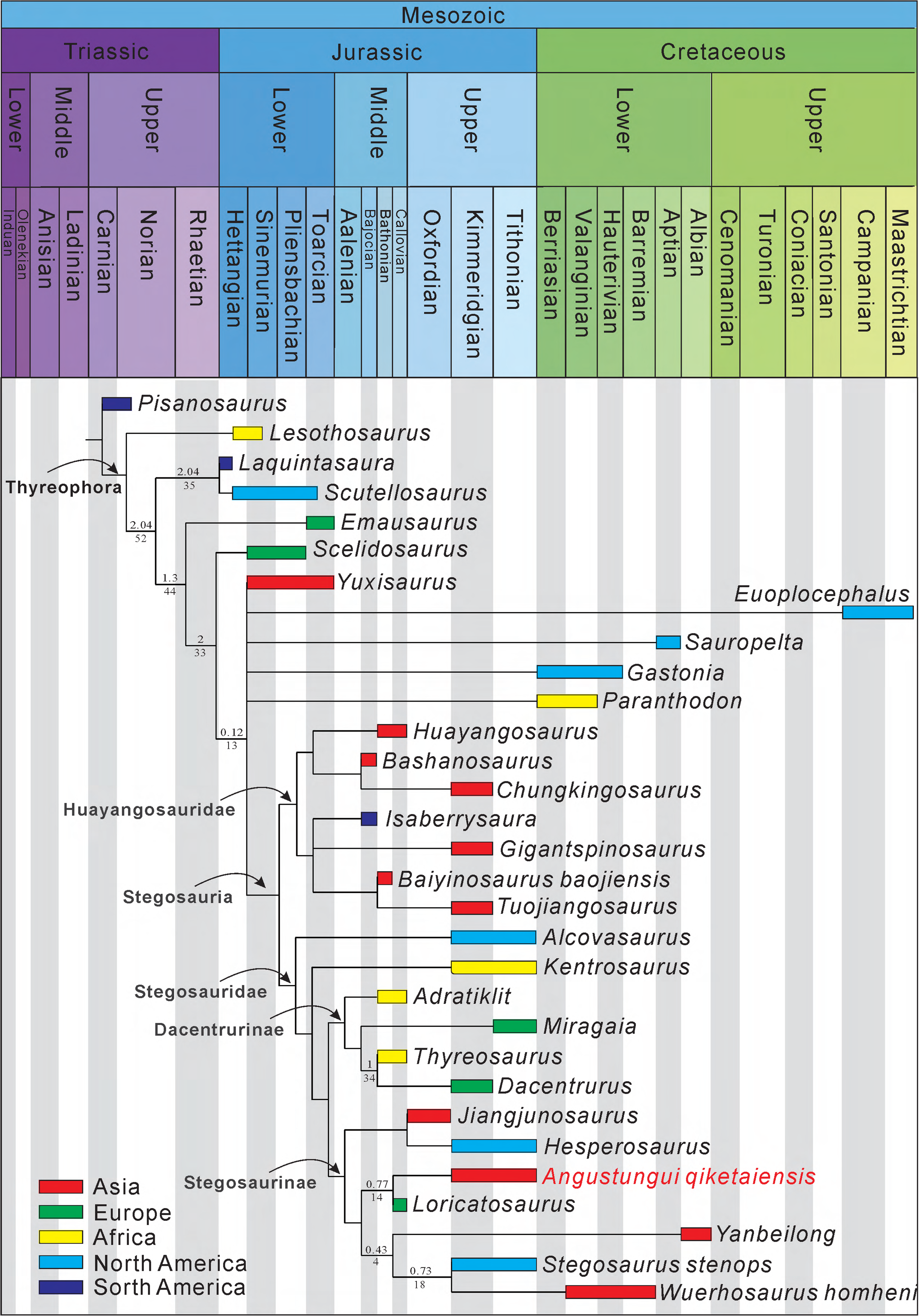
Phylogeny of stegosaurs is based on the strict consensus of the seven most parsimonious trees recovered by the phylogenetic analysis. Bremer support and bootstrap support percentages are indicated above and below the line.

## DISCUSSION

### Taxonomic remarks

*Angustungui* differs from other stegosaurs in bearing several autapomorphies. The anterior centroparapophyseal lamina (ACPL) of the dorsal vertebrae is drawn into the anterolateral margin of the centrum, unlike *Stegosaurus* and *Bashanosaurus* ACPL merges ventrally with the neural arch before reaching the neurocentral suture (Dai et al. 2022; Maidment et al. 2015), and *Adratiklit* that ACPL is drawn into anteriorly projecting rugosities on either side of the neural canal (Maidment et al. 2020). In the ilium, the preacetabular process projects approximately parallel to the parasagittal plane in dorsal or ventral view and lies approximately horizontal in lateral view, and a ventromedial flange backs the acetabulum. In other stegosaurs, the ventromedial flange backing the acetabulum is present only in *Kentrosaurus*, but the anterior iliac process diverges widely from the parasagittal plane in *Kentrosaurus* (Galton 1982). The ungual phalanx is claw-shaped, different from other stegosaurs with hoof-shaped ungual. In stegosaurs, the parascapular spine is only preserved in a few taxa, including *Loricatosaurus*, *Kentrosaurus* and *Gigantspinosaurus* (Galton 1982, 1985; Hao et al. 2018). The basal plate of the parascapular spine of *Angustungui* is sub-triangular in outline, different from *Loricatosaurus* with a sub-quadrangular basal plate and *Kentrosaurus* with a sub-circular basal plate (Galton 1982, 1985). In addition, the base of the spine of the parascapular spine length is more than one-third of the lateral margin of the basal plate length, which is more robust than in *Gigantspinosaurus* (Hao et al. 2018).

*Jiangjunosaurus* and *Wuerhosaurus homheni* are also from the Xinjiang Uygur Autonomous Region, China. In addition to the autapomorphies, *Angustungui* also has some differences with them. In *Angustungui*, the lateral surface of the cervical centra is deeply concave, but the lateral surface of the centrum of the cervical vertebra has openings in *Jiangjunosaurus* (Jia et al. 2007). The ventral keel of the cervical centrum is absent in *Angustungui*, but it is present in *Jiangjunosaurus* (IVPP V14724). The pedicels of the dorsal vertebrae are short dorsoventrally in *Angustungui*, unlike *Wuerhosaurus homheni* (IVPP 4006), which is greatly elongated. The neural arches of the dorsal vertebrae of *Wuerhosaurus homheni* (IVPP 4006) are deeply excavated dorsal to the neural canal in anterior view (Maidment et al. 2008), but it is absent in *Angustungui*. Maidment et al. 2008 (2008) referred to *Wuerhosaurus homheni* as a new species of *Stegosaurus*, *S*. *homheni*. *Wuerhosaurus homheni* was originally diagnosed by the following autapomorphies: the dorsal neural canals are small and not enlarged; the anterior processes of the ilia are separated by a wide-angle; the bony plates are large, long, and rather low in profile (Dong 1990, 1993). In dorsal vertebrae, the ratio of the neural arch to neural canal height and the ratio of the centrum height to the neural canal is close between *Wuerhosaurus homheni* and *Stegosaurus*. The anterior processes of the ilia are also separated by wide angle in *Stegosaurus* (Maidment et al. 2015). It is unknown whether the dorsal part of the plate of *Wuerhosaurus homheni* is complete because the current location of the plate is unknown. In addition, the plate of *Angustungui* and plate 6 of *Stegosaurus* are also longer than they are tall (Maidment et al. 2015). Although *Wuerhosaurus homheni* cannot be diagnosed based on these characteristics, there are also some differences with *Stegosaurus stenops*: the neural arches of the dorsal vertebrae of *Wuerhosaurus homheni* (IVPP 4006) are deeply excavated dorsal to the neural canal in anterior view; the dorsosacral vertebrae ribs of *Wuerhosaurus homheni* fuse to dorsal margins of the first true sacral vertebra, but the dorsosacral vertebrae ribs of *Stegosaurus stenops* fuse to the medial margin of preacetabular process of the ilium; the coracoid of *Wuerhosaurus homheni* is anteroposteriorly longer than dorsoventrally high in lateral view, but the coracoid of *Stegosaurus stenops* is sub-circular in outline (Maidment et al. 2015). In particular, the deeply excavated dorsal to the neural canal of the dorsal vertebrae is also present in *Yanbeilong* (Jia et al. 2024). So, we recover the Early Cretaceous Asian *Wuerhosaurus homheni* as a separate genus from the Jurassic American genus *Stegosaurus*. The revised diagnosis of *Wuerhosaurus homheni* is: that the neural arches of the dorsal vertebrae are deeply anteriorly excavated dorsal to the neural canal.

### Biogeographical insights

*Angustungui* has many morphologies similar to stegosaurs from Europe, Africa and North America, especially Europe. The lateral surfaces of the centra of the cervical vertebrae and dorsal vertebrae are deeply concave in *Angustungui*, similar to *Dacentrurus* and *Loricatosaurus* from Europe (Galton 1985, 1991), *Adratiklit* and *Thyreosaurus* from Africa (Maidment et al. 2020; Zafaty et al. 2024) and different from Asian taxa. The prezygapophysis of the dorsal vertebrae presents a distinct dorsal process in *Angustungui*, which is also present in *Dacentrurus* and *Adratiklit* (Maidment et al. 2020; Sánchez-Fenollosa et al. 2024), but absent in Asian taxa. There is a distinct fossa on the posterolateral surface of the prezygapophysis of the *Angustungui* dorsal vertebrae, also present in *Loricatosaurus* (BMNHR3167), *Craterosaurus* from Europe (Galton 1981) and *Kentrosaurus* (MB R.1930) from Africa (Maidment et al. 2008), but absent in Asian taxa. The dorsal process the transverse process of the anterior caudal vertebrae is present in *Angustungui*, which is also present in *Dacentrurus*, *Loricatosaurus*, and the North American taxa *Stegosaurus*, *Hesperosaurus* and *Alcovasaurus* (Carpenter et al. 2001; Galton 1985; Gilmore 1914; Sánchez-Fenollosa et al. 2024), but absent in Asian taxa. Furthermore, *Angustungui* is recovered as the sister taxon of *Loricatosaurus* in our phylogenetic analysis based on two synapomorphies (69 and 114). Therefore, *Angustungui* is a representative taxon of Stegosauridae from Asia, together with the Oxfordian *Jiangjunosaurus* indicating that Stegosauridae dispersed to East Asia before or during the Oxfordian. The discovery of *Angustungui* adds credibility to the idea that the isolation of East Asia, and associated endemism, might not have been as profound as previously supposed (Mannion et al. 2019; Xu et al. 2018).

### The evidence of the stegosaurian claw-shaped unguals

The stegosaurs and ankylosaurs constitute the ‘broad-footed’ thyreophorans (Eurypoda) (Sereno 1999). All preserved ungual phalanges in ankylosaurs and stegosaurs are broad hoof-shaped (Raven et al. 2023). Although the non-terminal phalanx of *Angustungui* is broad as in other stegosaurs, the ungual phalanx of *Angustungui* is narrow and claw-shaped, unlike other stegosaurs with broad and hoof-shaped ungual phalanx. On the one hand, this is interpreted by the fact that only a few well-preserved stegosaurian specimens have been found. In particular, among stegosaurs, the unguals are only preserved in *Huayangosaurus*, *Kentrosaurus*, *Gigantspinosaurus*, *Wuerhosaurus homheni*, *Stegosaurus stenops* and *Angustungui* (Galton 1982; Hao et al. 2018; Maidment et al. 2006, 2008, 2015). *Angustungui* may have retained a plesiomorphic condition perhaps due to the special living environment. The claw-shaped ungual of *Angustungui* is more similar to the basally branching thyreophoran *Scelidosaurus* (Norman 2020), which was used for weight support and digging. However, some morphologies of the pelvic girdle of *Angustungui* are also similar to *Scelidosaurus* (Norman 2020): in the ilium, the preacetabular process projects roughly parallel to the parasagittal plane in dorsal or ventral view and lies approximately horizontal in lateral view, and a ventromedial flange backing the acetabulum; the pubic shaft is much more slender compared to the prepubis and the shaft of the ischium. In short, *Angustungui* provided evidence that some Eurypoda, at least stegosaurs, had claw-shaped ungual phalanges.

## CONCLUSIONS

A partial skeleton collected from the Upper Jurassic Qigu Formation of Xinjiang, China, represents a new genus and species of stegosaur dinosaur, which we name *Angustungui qiketaiensis*. It can be distinguished from all other stegosaurs by several autapomorphies of the dorsal vertebrae, ilium, ungual and parascapular spine. *Angustungui* recovered as the sister taxa of *Loricatosaurus* in our phylogenetic analysis, and it is a representative taxon of Stegosauridae from Asia. The claw-shaped ungual of *Angustungui* provides evidence that Eurypoda, at least stegosaur, has claw-shaped unguals.

## ACKNOWLEDGEMENTS

We thank Jiang S. and Hao B.-Q. from the Zigong Dinosaur Museum, Gao B.-C. from the Chongqing Natural History Museum and Zheng F. from the Institute of Vertebrate Paleontology and Paleoanthropology for access to specimens in their care. We thank the Shanshan Bureau of Natural Resources, Turpan City, Xinjiang Uygur Autonomous Region, China for supporting this research. We also appreciate the thoughtful reviews provided by anonymous reviewers and the editors who contributed to enhancing an earlier version of this manuscript.

## AUTHOR CONTRIBUTIONS

Li Ning wrote the main manuscript text. Chen Guozhong prepared all the figures and measured the specimens. Octávio Mateus improved the manuscript text and helped with English language editing. Jiang Tao, Li Daqing, You Hailu and Peng Guangzhao review the manuscript text. Xie Yan and Li Daqing oversaw the project. All authors reviewed the manuscript.

## SUPPLEMENTARY DATA

Measurements of the holotype (SS V16001) and paratype (SS V16002) can be found in Supplementary 1. The revised character list and data matrix can be found in supplementary 2 and 3.

## DATA AVAILABILITY

All data generated or analysed during this study are included in this published article and its supplementary information.

